# Isolation and sequencing of AGO-bound RNAs reveals characteristics of mammalian stem-loop processing *in vivo*

**DOI:** 10.1101/294488

**Authors:** Ian M. Silverman, Sager J. Gosai, Nicholas Vrettos, Shawn W. Foley, Nathan D. Berkowitz, Zissimos Mourelatos, Brian D. Gregory

**Author notes:** To whom correspondence should be addressed. Brian D. Gregory, Department of Biology, University of Pennsylvania, 433 S. University Ave., Philadelphia, PA 19104. These authors contributed equally to this work (Ian M. Silverman, Sager J. Gosai). Authors Note*: We have deposited raw and processed sequencing data to GEO under the accession GSE71710, which is currently available to reviewers at http://www.ncbi.nlm.nih.gov/geo/query/acc.cgi?token=mbursywspvkjxoh&acc=GSE71710. We have also created a website to house coverage profiles and RNAfold diagrams generated in this study at http://gregorylab.bio.upenn.edu/AGO_IP_Seq/.

## Abstract

MicroRNA precursors (pre-miRNAs) are short hairpin RNAs that are rapidly processed into mature microRNAs (miRNAs) in the cytoplasm. Due to their low abundance in cells, sequencing-based studies of pre-miRNAs have been limited. We successfully enriched for and deep sequenced pre-miRNAs in human cells by capturing these RNAs during their interaction with Argonaute (AGO) proteins. Using this approach, we detected > 350 pre-miRNAs in human cells and > 250 pre-miRNAs in a reanalysis of a similar study in mouse cells. We uncovered widespread trimming and non-templated additions to the 3’ ends of pre- and mature miRNAs. Additionally, we created an index for microRNA precursor processing efficiency. This analysis revealed a subset of pre-miRNAs that produce low levels of mature miRNAs despite abundant precursors, including an annotated miRNA in the 5’ UTR of the DiGeorge syndrome critical region 8 mRNA transcript. This led us to search for AGO-associated stem-loops originating from other mRNA species, which identified hundreds of putative pre-miRNAs derived from human and mouse mRNAs. In summary, we provide a wealth of information on mammalian pre-miRNAs, and identify novel microRNA and microRNA-like elements localized in mRNAs.

## INTRODUCTION

MicroRNAs (miRNAs) are ~22 nucleotide (nt) small RNAs (smRNAs) that negatively regulate gene expression at the post-transcriptional level by repressing translation and/or promoting degradation of target messenger RNAs (mRNAs) (1,2). Animal miRNAs are generated in a two-step process, whereby miRNA precursors (pre-miRNAs) are first cleaved from their primary transcripts (pri-miRNAs) by the action of the DGCR8/DROSHA microprocessor complex (3,4). Alternative pre-miRNA biogenesis pathways have been described that bypass the microprocessor. For example, mirtron loci generate pre-miRNAs in a splicing-dependent and DROSHA-independent fashion (5,6). Pre-miRNAs are then transported into the cytoplasm by Exportin-5 for further processing (7,8).

Cytoplasmic pre-miRNAs are matured by the miRNA loading complex (miRLC) which is composed of the type III endonuclease DICER, the double-stranded RNA-binding protein TRBP, and the miRNA effector protein Argonaute (AGO) (9–11). DICER cleaves the pre-miRNA to reveal a ~22 nt miRNA duplex, consisting of the upstream miRNA (denoted as the 5p miRNA) and downstream miRNA (denoted as the 3p miRNA). One of these strands is selectively loaded into AGO to form the RNA-induced silencing complex (RISC). Alternatively, AGO2 has been shown to directly cleave several pre-miRNAs, which are then further processed by the poly(A)-specific ribonuclease (PARN) to give rise to mature miRNAs. This class of miRNAs is known as the AGO2-cleaved pre-miRNAs (ac-pre-miRNAs) (12–15). Such DICER-independent processing involves the pre-miRNA deposit complex (miPDC), which is composed of just AGO2 and a pre-miRNA (16).

The miRNA biogenesis mechanism is further complicated by the fact that both pre- and mature miRNAs can be post-transcriptionally modified at the 3’ end by trimming or by non-templated addition of ribonucleotides, especially uridine and adenine (17–24). In general, mono-uridylation is thought to re-establish a 3’ 2 nt overhang, which is required for efficient DICER cleavage. In contrast oligo-uridylation has been shown to be a signal for degradation and usually occurs after AGO2-mediated slicing of pre-miRNAs. However, most detailed studies of pre-miRNAs have been performed on a small number of pre-miRNA sequences, thus our understanding of the overall landscape, and global function of these modifications remains elusive.

High-throughput sequencing of total or AGO-bound smRNAs in numerous cells, tissues, and organisms has provided a wealth of information about miRNAs. In concert with bioinformatic approaches, these datasets have been expanded to identify thousands of novel miRNAs (25,26). In contrast, sequencing of pre-miRNAs has been challenging, due to the presence of other RNA species, including the highly abundant transfer RNAs (tRNAs) and small nucleolar RNAs (snoRNAs), that exist in the same size range as pre-miRNAs. Attempts to use size selection to sequence pre-miRNAs have achieved < 1% of total sequenced clones corresponding to pre-miRNAs even after selective depletion of abundant species (27). Alternatively, primer-based approaches have been applied, but these methods restrict analysis to known miRNA species and thus cannot be used for discovery (20,21,28). Thus, there is a need for unbiased, sequencing based approaches to gain a more comprehensive understanding of pre-miRNA expression and sequence content.

Leveraging the knowledge that AGO is an integral component of the pre-miRNA processing complexes (miRLC and miPDC), we recently developed an approach to enrich for pre-miRNAs by immunoprecipitating AGO proteins and subsequently cloning RNAs from 50-80 nucleotides (nts). Using this approach, pre-miRNA libraries from mouse embryonic fibroblasts (MEFs) were generated with > 40% of reads mapping to miRNA loci (19). Thus, this strategy can be used to efficiently study pre-miRNAs globally without the need for primer-based approaches or depletion of abundant RNA species. Here, we applied this approach to isolate and sequence pre-miRNAs and mature miRNAs from human embryonic kidney (HEK293T) cells. We developed a bioinformatic pipeline to capture post-transcriptional modifications, which we applied to data generated in this study from HEK293T cells, as well as to the previous data generated in MEFs. Our results provide global insights into pre-miRNA processing and provide an alternative strategy for identifying pre-miRNAs and other AGO-associated stem-loops in transcriptomes of interest.

## MATERIALS AND METHODS

### Cell culture

HEK293T cells were grown to 70-80% confluence in 15 cm tissue culture plates with DMEM media supplemented with 10% FBS and 1X pen/strep at 37°C and 5% CO_2_.

### AGO-IP-sequencing

Pre-miRNA-seq and miRNA-seq preparation was based on previous work (19,29,30). HEK293T cells were lysed in RSB 200 (20 mM Tris-HCl pH 7.4, 200 mM NaCl, 2.5 mM MgCl_2_, 0.5% NP-40) supplemented with 0.2 U/μl RNasein (Promega) and protease inhibitor cocktail (Roche). Cleared lysates were incubated with Protein G agarose beads (Life Technologies) conjugated to AGO 2A8 antibody or non-immune serum (NMS), for 1.5 h at 4°C. After four 5 min washes in lysis buffer, a sample of beads was drawn for western blot analysis while the rest was subjected to RNA extraction with Trizol (Life Technologies). RNA was dephosphorylated with antarctic phosphatase (NEB) and P^32^-γ-ATP labeled with T4 polynucleotide kinase (NEB).

5’ end radiolabeled RNAs were resolved on a 7 M urea, 15% polyacrylamide gel. Gel slices from 20-25 nt (miRNA-seq) and 50-80 nt (pre-miRNA-seq) were recovered and ligated to miRCat 3’ Linker-1 (IDT) with T4 RNA Ligase 2 Truncated (NEB), in the presence of 15% PEG8000. Ligation products were PAGE purified and reverse transcribed by a 5’ phosphorylated RT primer containing 3’ and 5’ adaptor complementary sequences. To do this, RT primer was mixed with RNA and heated at 80°C for 5 min, then incubated at 60°C for 5 min and finally cooled down to room temperature before the remaining reagents were added (Affinityscipt, Stratagene) (31). cDNAs were resolved and purified from a 7 M urea, 6% polyacrylamide gel, then circularized with CircLigase I (Epicentre) and PCR amplified with Phusion DNA polymerase (NEB). The amplicon yield was boosted with a second PCR amplification round, then size selected on a 3% metaphor gel (Lonza), purified, and sent for deep sequencing at the Next-Generation Sequencing Core at the University of Pennsylvania. All oligonucleotide sequences used in this study can be found in the attached document (**Table S9**).

### Mapping Pipeline

The first 20 nt of the 3’ adapter sequence *CTGTAGGCACCATCAATAGA* was used to trim adapter sequence from the raw reads using cutadapt (v1.4.2). Identical reads were collapsed but clone information was retained to reduce computational time. Trimmed reads were sequentially mapped to miRBase (v20) and RefSeq annotated spliced transcript models (hg19 or mm10, downloaded on 06082015), with a two-stage alignment strategy with Bowtie2 and EMBOSS-WATER. First, Bowtie2 was used to map the 5’ regions of reads to either miRbase (v20) primary miRNA sequences with a 50 nt extension on the 3’ end or RefSeq annotated spliced transcript models. For pre-miRNA-seq, the first 35 nt were used in the initial alignment step, whereas 18 nt were used for miRNA- and smRNA-seq. Following Bowtie2 alignment, reads were extended by local alignment with EMBOSS-WATER (with parameters: -gapopen 10.0 -gapextend 0.5). Multimapped reads were partially resolved by selecting the longest, highest scoring alignments. Mismatches detected at the 3’ ends of reads were considered as non-templated additions and analyzed separately. We filtered unmapped reads from pre-miRNA-seq for known rRNA, snoRNA, snRNA, tRNA, mitochondrial transcripts, and repeat-masked sequences before mapping to RefSeq.

### Analysis of miRNA trimming and non-templated tailing

We tabulated non-templated additions revealed by our mapping pipeline for reads, which mapped to high-confidence miRBase annotations. Additionally, we calculated templated-extension and trimming by comparing mapped ends of pre-miRNA-seq, miRNA-seq, and smRNA-seq reads against annotated 5p and 3p miRNA ends.

### Identification of AGO-2 cleaved pre-miRNAs

For each miRBase miRNA, we calculated the percentage of trimmed pre-miRNA-seq reads that ended between 8-15 nt upstream of the 3’ end of the 3p miRNA or downstream of the 5’ end of the 5p miRNA. We identified pre-miRNAs with >1% of such cleavage events in the 3p arm as putative ac-pre-miRNAs. For cleavage events in the 5p arm, we took a more conservative approach, requiring 5% of reads to terminate in this region and > 50% of mature miRNA-seq reads in the 3p arm compared to 5p arm.

### Determination of MPAR

We calculated the mature to precursor abundance ratio (MPAR) for miRBase miRNAs or AGO-associated stem-loops as the generalized log ratio (glog) (32–34) of miRNA-seq (smRNA-seq) to pre-miRNA-seq RPM coverage (nmi, npr, respectively) as follows:

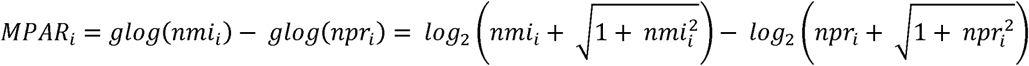

Thus MPAR can be calculated for loci that are not represented in either the pre-miR-seq or miR-seq library. Loci with no coverage in either sequencing library were omitted from this analysis.

### Identification of AGO-associated stem-loops derived from mRNAs

Pre-miRNA-seq reads that failed to map to miRBase and were not removed by mapping to known rRNA, snoRNA, snRNA, tRNA, mitochondrial transcripts, and repeat-masked sequences were aligned to RefSeq transcripts with Bowtie2. After filtering for best matches for reads with less than 4 mismatches, we used Pycioclip to call significant peaks with a modified False Discovery Rate < 0.01 (35). To remove abundant genes with high numbers of mappings but no local peaks, we filtered out peaks that were greater than 200 nt in length. We then chose the most abundant clone and predicted its secondary structure with RNAfold using standard parameters (36). We again filtered clusters, requiring that they had greater than 15 bp in the longest hairpin and a total MFE of less than −0.3 kcal/mol/nt. We analyzed non-templated tailing for reads mapping to these hairpins as described above for our miRNA analysis.

### icSHAPE analysis

*In vivo* icSHAPE data from mouse ES cells was downloaded from GEO using the accession number GSE60034. Only scores in the top or bottom 10^th^ percentile were used for analysis. icSHAPE reactivity scores were overlaid on RNAfold diagrams using RNAplot (36).

### Small RNA Northern blotting

20-30 μg of total RNA was resolved on 8 M urea 12.5% PAGE and transferred to Hybond-N+ membranes (GE Healthcare). Blotted RNA was hybridized with γ-^32^P-ATP labeled LNA oligonucleotides (**Table S9**) in UltraHyb-Oligo Buffer (Ambion) at 50ºC overnight. Membranes were washed once in 2x SSC, 0.1% SDS at 50ºC and once in 0.1x SSC, 0.1% SDS at 50ºC before exposing to autoradiography film for 1 week at −80ºC.

### siRNA-mediated knockdown of DGCR8 and DROSHA

ON-TARGET plus siRNAs against Human *DGCR8 and DROSHA* were obtained from Dharmacon (J-015713-05-0002, J-015713-06-0002, J-016996-05-0002, J-016996-06-0002). siRNAs against Luciferase were generated by IDT. We performed two sequential siRNA transfections 48 hours apart with 45 pmol of siRNAs (22.5 pmol siRNA-1 and 22.5 pmol siRNA-2), 125 μL Opti-MEM (Life Technologies), and 6 μL Lipofectamine 2000 (Life Technologies) per reaction. Reactions were incubated at room temperature for 20 minutes. During this incubation, we seeded 6.0×10^5^ HEK293T cells per replicate in two wells of a 6-well plate (3.0×10^5^ cells/well) in 2 mL media. We then added the siRNA mixture to each well dropwise, and allowed cells to incubate at 37ºC in 5% CO_2_ for 48 hours. After 48 hours we repeated the transfection, harvesting the two wells per replicate (~1.2×10^5^ cells) and dividing them into four wells in a 6-well plate. We treated each well with 45 pmol of siRNAs, and allowed them to incubate at 37ºC in 5% CO_2_ for another 48 hours. Cells were then pooled and washed with PBS prior to storage at −80ºC.

### mRNA-seq

mRNA-seq was performed as previously described (37). Briefly, total RNA was purified from the HEK293T cell cultures (miRNeasy; Qiagen). Poly(A)^+^ RNA was isolated using oligo(dT) beads (Life Technologies). RNA was fragmented for 7 minutes using Fragmentation Reagent (Life Technologies). mRNA-seq libraries were then generated using the Illumina mRNA-seq kit (Illumina). Reads were trimmed with Cutadapt, mapped with Tophat2, and gene expression was quantified using HTseq (38–40). DEseq2 was used to perform differential expression analysis (41)

## RESULTS

### Isolation and sequencing of pre-miRNAs

To isolate and sequence pre-miRNAs we first immunoprecipitated AGO proteins from HEK293T cells with the pan-AGO-2A8 antibody (42) (Fig. 1A). RNA was purified from AGO immunoprecipitates, dephosphorylated and labeled with P^32^-γ-ATP. Autoradiography of the RNA gel showed a major band at 20-25 nt, representing mature miRNAs, and several other prominent bands between 50-80 nt, corresponding to the size range of pre-miRNAs (Fig. 1B). We excised gel slices from both of these regions and generated high-throughput sequencing libraries, referred to herein as miRNA-seq and pre-miRNA-seq libraries, respectively. Previous attempts to sequence pre-miRNAs have been unsuccessful due to the inaccessibility of the pre-miRNA 5’ ends. Therefore, we used a method that attaches the 5’ linker through a CircLigase-mediated cDNA circularization step (see **Methods**) (19,29).

**Figure 1.**
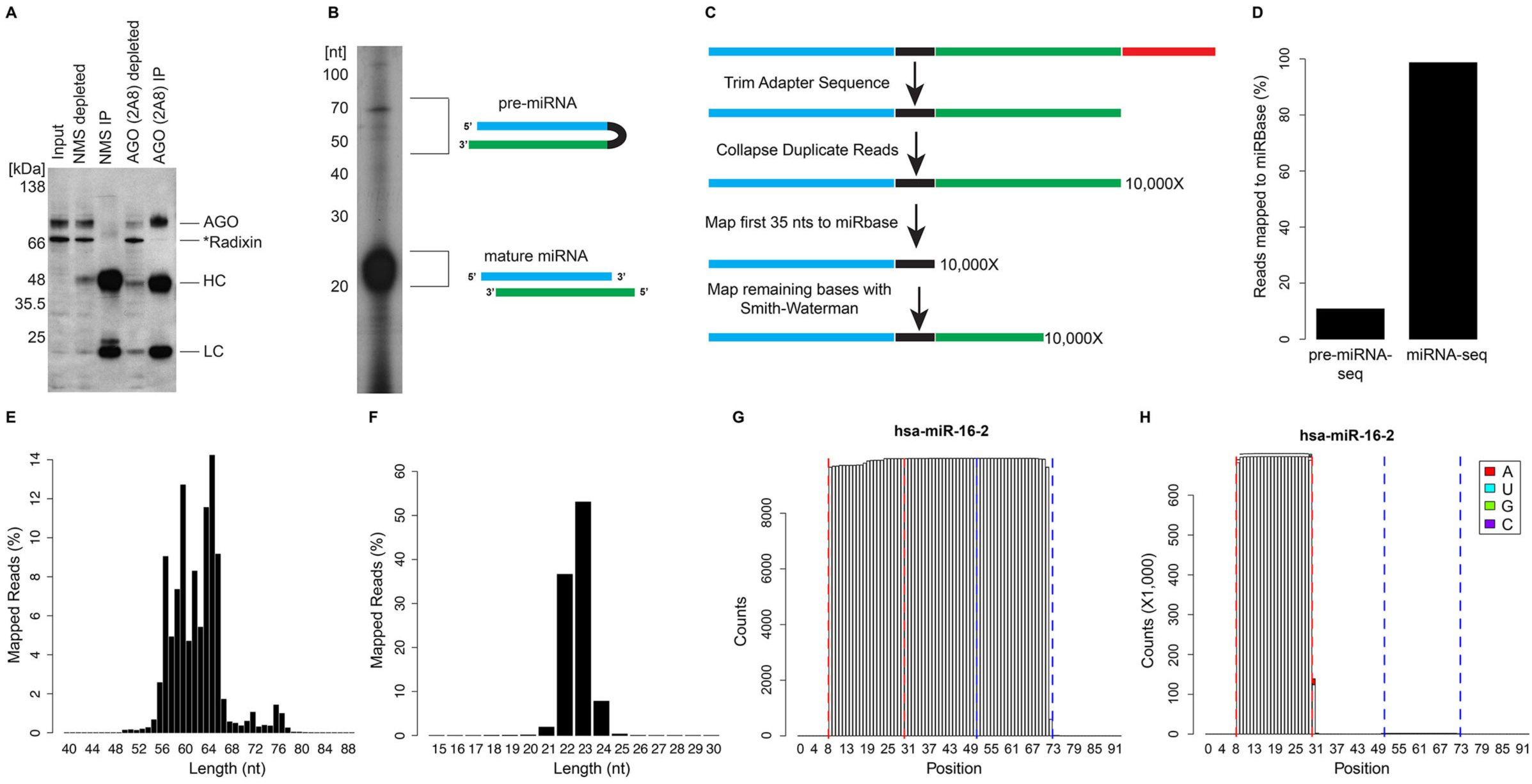
Isolation and sequencing of AGO interacting pre-miRNAs and mature miRNAs. A) Western blot of AGO-IP from HEK293T cells. AGO-2A8 and non-immune mouse serum (NMS) were used for immunoprecipitation. AGO-2A8 antibody was used for detection. *Radixin is known to cross-react with the 2A8 antibody. Immunoglobulin heavy chain (HC) and light chain (LC) are denoted. B) Autoradiography of RNA co-immunoprecipitated with AGO. Pre-miRNAs were excised from 50-80 nt size range and mature miRNAs were excised from 20-25 nt size range. C) Bioinformatics pipeline: Adapter sequences were trimmed and PCR duplicates were collapsed for efficiency. The first 35 nt of pre-miRNAs (18 nt of mature miRNAs) were mapped to miRBase (v20). Remaining bases were mapped with a Smith-Waterman aligner. D) Percent of reads mapping to human miRBase for HEK293T pre-miRNA-seq and miRNA-seq. E-F) Size distribution of pre-miRNA-seq (E) and miRNA-seq (F) reads mapping to miRBase. G-H) Coverage plot of pre-miRNA-seq (G) and miRNA-seq (H) reads mapping to hsa-miR-16-2 locus. White bars indicate templated nucleotides. Colored bars indicate non-templated additions. Dashed red and blue lines indicate boundaries of annotated 5p (blue) and 3p (red) mature miRNAs.

Using this approach, we successfully generated high-throughput sequencing libraries for both pre-miRNAs and mature miRNAs. We obtained 27.8 and 9.8 million reads with sufficient adapter sequence from the pre-miRNA-seq and mature miRNA-seq libraries, respectively (**Table S1**). Given that pre-miRNAs and mature miRNAs contain non-templated modifications at their 3’ ends, we reasoned that standard alignment pipelines would be limited in their ability to align these sequences to miRBase. Therefore, we developed an alignment pipeline that utilized the first 35 nt of the pre-miRNA-seq (18 nt for miRNA-seq) reads for alignment (Fig. 1C), followed by Smith-Waterman local alignment to extend the read as far as possible along the mapped miRNA sequence (see **Methods**) (43–45). After all possible nucleotide matches were made, we selected alignments with the lowest mismatch rate and captured non-templated modifications at the 3’ end.

We performed this analysis on pre-miRNA-seq, miRNA-seq, and smRNA-seq (without IP) from human HEK293T cells as well as pre-miRNA-seq and mature miRNA-seq data previously generated using the same technique on MEFs (19). We successfully aligned 10.8% of pre-miRNA-seq, 98.7% of miRNA-seq, and 52.4% of smRNA-seq reads to the human miRBase v20 annotation set (Figs. 1D, **S1A, and Table S1**). We were not surprised to find such high rates of mapping for mature miRNAs, however, a 10% mapping rate for pre-miRNAs is significantly higher than previous attempts to sequence pre-miRNAs without pre-miRNA specific primers (0.8%) (27). Using the datasets previously generated from MEFs, we had a much higher rate of pre-miRNA-seq reads mapping (44%), but mapped fewer miRNA-seq reads (90.3%) (Fig. S2A **and Table S1**). These differences likely represent variable experimental conditions and amounts of starting material and/or biological differences in the smRNA populations between these two mammals.

Overall, we obtained pre-miRNA-seq reads mapping to 367 annotated human microRNAs and 267 annotated mouse microRNAs (**Table S2**). For libraries prepared from smaller RNA species, we mapped reads to 931 (HEK239T miRNA-seq), 567 (MEF miRNAs-seq), and 1,364 (HEK293T smRNA-seq) miRBase miRNAs. We examined the size distribution of mapped pre-miRNA-seq reads and found that they were broadly distributed between 55-65 nt in both cell types (Figs. 1E and S2B). In contrast, miRNA-seq reads, were tightly distributed between 21-24 nt, in both HEK293T AGO-bound and total cellular fractions, as well as in MEFs (Figs. 1F, S1B, and S2C). We determined the abundance, end concordance, and non-templated additions for all mapped miRNAs and generated coverage plots to represent these data, which are available for download at http://gregorylab.bio.upenn.edu/AGO_IP_Seq/ (**Table S2; examples in** Figs. 1G-H). Together, our biochemical and bioinformatic approaches provide a data rich resource for the global and unbiased analysis of pre-miRNAs in two mammals.

### Diverse ends of AGO-bound pre-miRNAs

It is well established that pre-miRNA trimming and non-templated tailing (uridylation) is a mechanism of regulation (17,18,20). However, most studies have investigated individual miRNAs or used targeted approaches to examine a predetermined subset of miRNAs. Using our novel datasets, we examined the end concordance of sequenced pre-miRNA-seq (Figs. 2A and S2D) and miRNA-seq reads (Figs. 2B and S2E), relative to high confidence human and mouse miRBase miRNA ends. We found that the majority (>90%) of 5’ read ends of pre-miRNAs coincided with annotated 5’ ends of 5p miRNAs for both human and mouse (Figs. 2A and S2D). In contrast 3’ read ends of pre-miRNAs were highly variable, with only 40% and 20% of reads ending precisely at the annotated 3’ end of 3p miRNAs in human and mouse cells, respectively (Figs. 2A and S2D). Relatedly, mature miRNAs (Figs. 2B and S2E) and smRNAs (Fig. S1C) displayed higher variation in their 3’ ends relative to 5’ ends for both 5p and 3p miRNAs, but not nearly to the extent that was observed for pre-miRNAs (Figs. 2A and S2D). Overall, these results reveal that 3’ end variation is more common in pre-miRNAs than in mature miRNAs.

**Figure 2.**
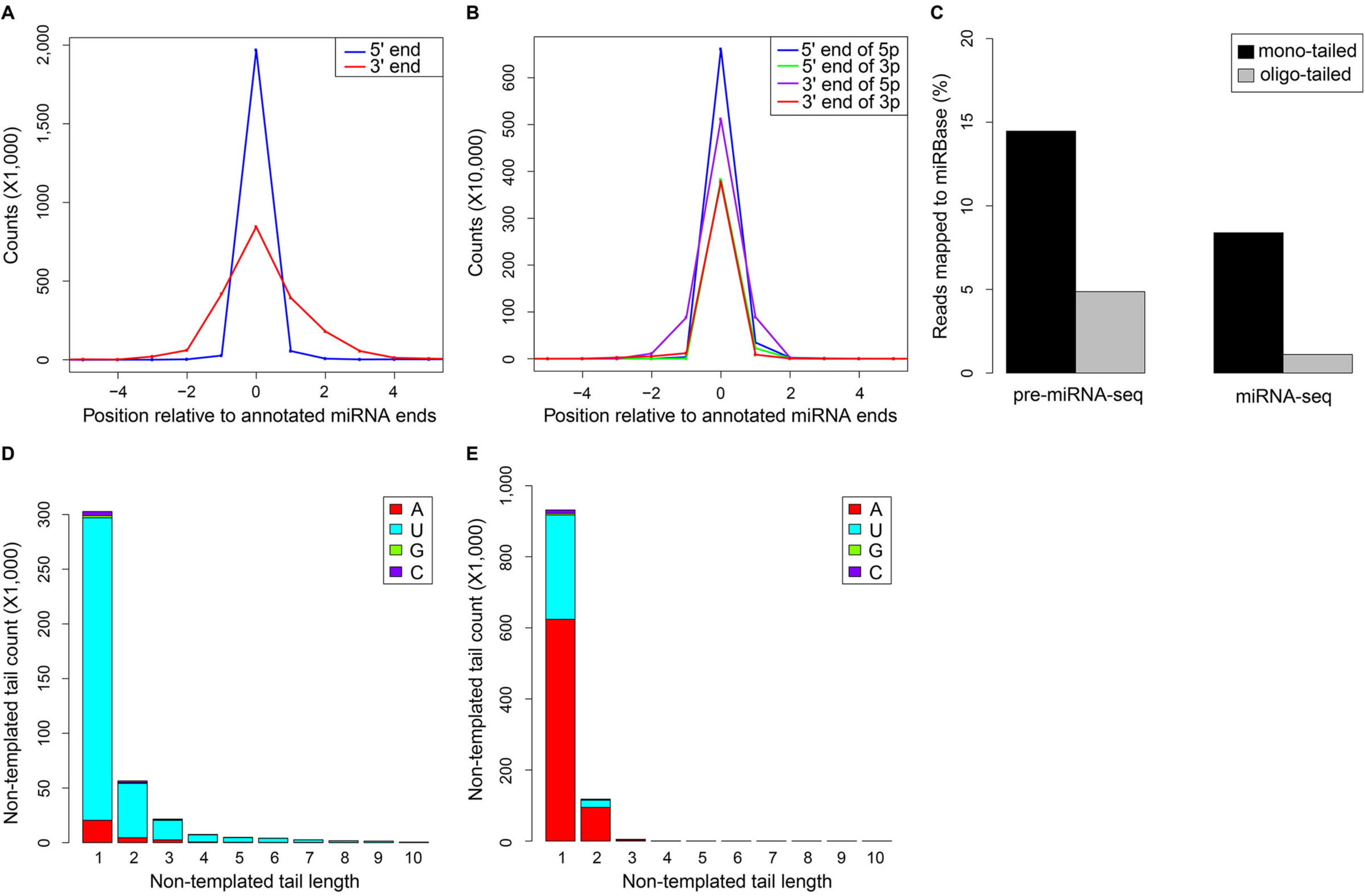
pre-miRNA-seq identifies diverse ends of pre-miRNAs. A) Distribution of pre-miRNA-seq read ends relative to annotated 5’ end of 5p and 3’ end of 3p miRNAs for high confidence human miRNAs. Positive and negative values indicate trimming and extension of reads, respectively. B) Distribution of miRNA-seq read ends relative to annotated ends of 5p and 3p miRNAs for high confidence human miRNAs. Positive and negative values indicate trimming and extension of reads, respectively. C) Percentage of reads with mono-tails and oligo-tails for HEK293T pre-miRNA-seq and miRNA-seq reads mapped to high confidence human miRBase miRNAs. D-E) Non-templated 3’ end tail length and sequence content for pre-miRNA-seq (D) and miRNA-seq (E) reads mapping to miRBase high confidence human miRNAs.

Pre-miRNAs and miRNAs are post-transcriptionally modified at their 3’ ends through the action of TUT4 and TUT7 terminal uriydyl transferases (TUTases) (17,18,46,47). We found that 14.5% of human and 17.4% of mouse pre-miRNAs contained single nucleotide additions to their 3’ ends, whereas 4.9% and 7.1% had more than two non-templated additions on their 3’ ends in human and mouse high confidence miRBase-annotated miRNAs, respectively (Figs. 2C and S2F). For mature miRNAs, we also found a large number of single nucleotide additions at the 3’ end; 8.4% of human and 12.5% of mouse mature miRNAs. However, tails greater than one nucleotide were much less frequent, with only 1.1% of human and 2.2% of mouse mature miRNAs containing long tails. The lower overall modification level of mature miRNAs is consistent with the hypothesis that oligo-uridylation is a signal for degradation, whereas mono-uridylation establishes the 3’ 2 nt overhang, required for efficient DICER cleavage.

For cellular smRNA-seq from HEK293T cells, we observed lower levels of single nucleotide additions (4.0%) and higher levels of multiple non-templated 3’ additions (3.3%), compared to AGO-bound miRNAs (8.4% and 1.1% respectively in human). (Fig. S1D). These differences may indicate that short oligo-nucleotide additions, which in some cases can promote pre-miRNA processing, could in fact prevent proper AGO loading of the 3’ mature miRNA. Collectively, these data demonstrate that non-templated additions are widespread in both pre-miRNAs and mature miRNAs, and that extended tails are more frequent in pre-miRNAs compared to mature miRNAs.

We also generated metaplots to analyze the sequence content of non-templated additions to the 3’ end of pre-miRNAs (Figs. 2D and S2G) and mature miRNAs (Figs. 2E, S1E, and S2H). Uridine (denoted as T) was by far the most common addition to human and mouse pre-miRNAs, and was even more prevalent in positions past the first non-templated nucleotide (Figs. 2D and S2G). For mature miRNAs, adenosine was the most common non-templated addition, with much lower levels of uridine as compared to pre-miRNAs (Figs 2E, S1E, and S2H). When examining non-templated additions in total smRNA-seq data, we found that 2 nucleotide tails were much more common than in AGO-bound miRNAs (Fig. S1E). Collectively, these data show that mono- and oligo-tailing are widespread in human and mouse pre- and mature miRNAs. Furthermore, they reveal that uridylation is more common in pre-miRNAs, especially after the first nucleotide, whereas adenylation is more common in mature miRNAs.

### Identification of AGO2-cleaved pre-miRNAs

AGO2, is unique amongst the mammalian AGO proteins in that it has slicing activity (48–50). In fact, AGO2 has been demonstrated to cleave a subset of pre-miRNAs, named AGO-cleaved pre-miRNAs (ac-pre-miRNAs) (13). The best characterized of the ac-pre-miRNAs, pre-miR-451, is trimmed by PARN following AGO2 cleavage to generate the mature miRNA sequence (12,14,15,51). However, the extent to which this process occurs in mammalian pre-miRNA populations has not been determined. AGO2 cleavage events are known to occur around 10 nucleotide upstream of the 3’ end of the pre-miRNA and give rise to 5p miRNAs (13). To search for these events, we calculated the percentage of pre-miRNA-seq reads that terminated 8-15 nucleotide from the 3’ end of the 3p miRNA for each miRNA. In mouse, for which ac-pre-miRNAs are best characterized, we observed all previously identified ac-pre-miRNAs with the exception of miR-451, which is not expressed in MEFs (Table 1). In our human dataset, we also observed previously identified ac-pre-miRNAs, including 9 members of the let-7 family and miR-9-2 (**Table S3**). From this approach, we identified 7 putative ac-pre-miRNA candidates in humans and 37 in mouse, including hsa-miR-455 and mmu-miR-335 (Figs. 3A-B and Tables 1 and S3).

**Figure 3.**
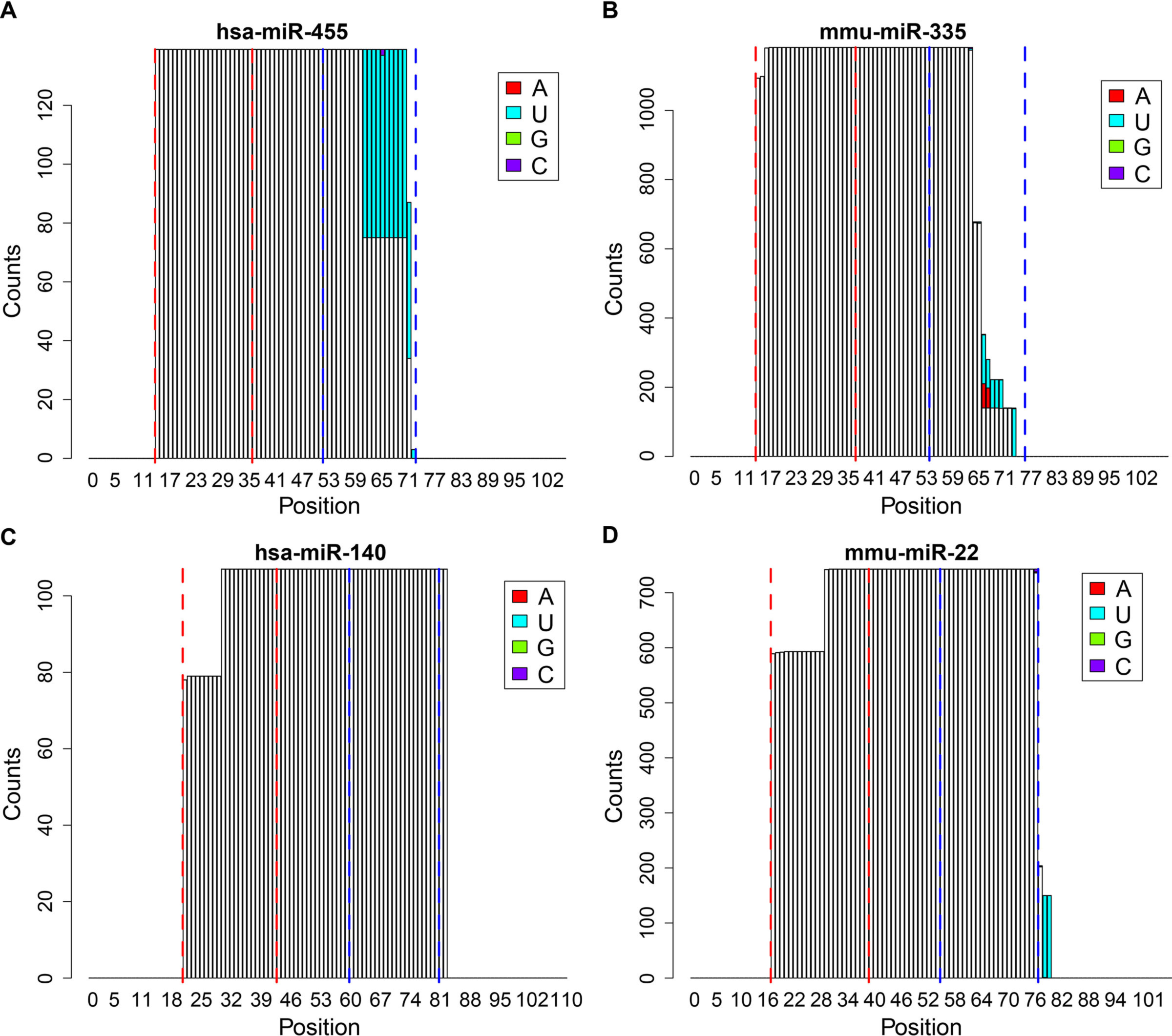
*De novo* identification of AGO2-cleaved pre-miRNAs. A-D) Examples of novel ac-pre-miRNAs identified from pre-miRNA-seq. hsa-miR-455 (A) and mmu-miR-335 (B) are novel ac-pre-miRNAs with cleavage in the 3p miRNA. hsa-miR-140 (C) and mmu-miR-22 (D) are novel ac-pre-miRNAs with cleavage in the 5p miRNA. White bars indicate templated nucleotides, colored bars indicate non-templated additions. Dashed red and blue lines indicate boundaries of annotated 5p (blue) and 3p (red) miRNAs.

**Table 1:**
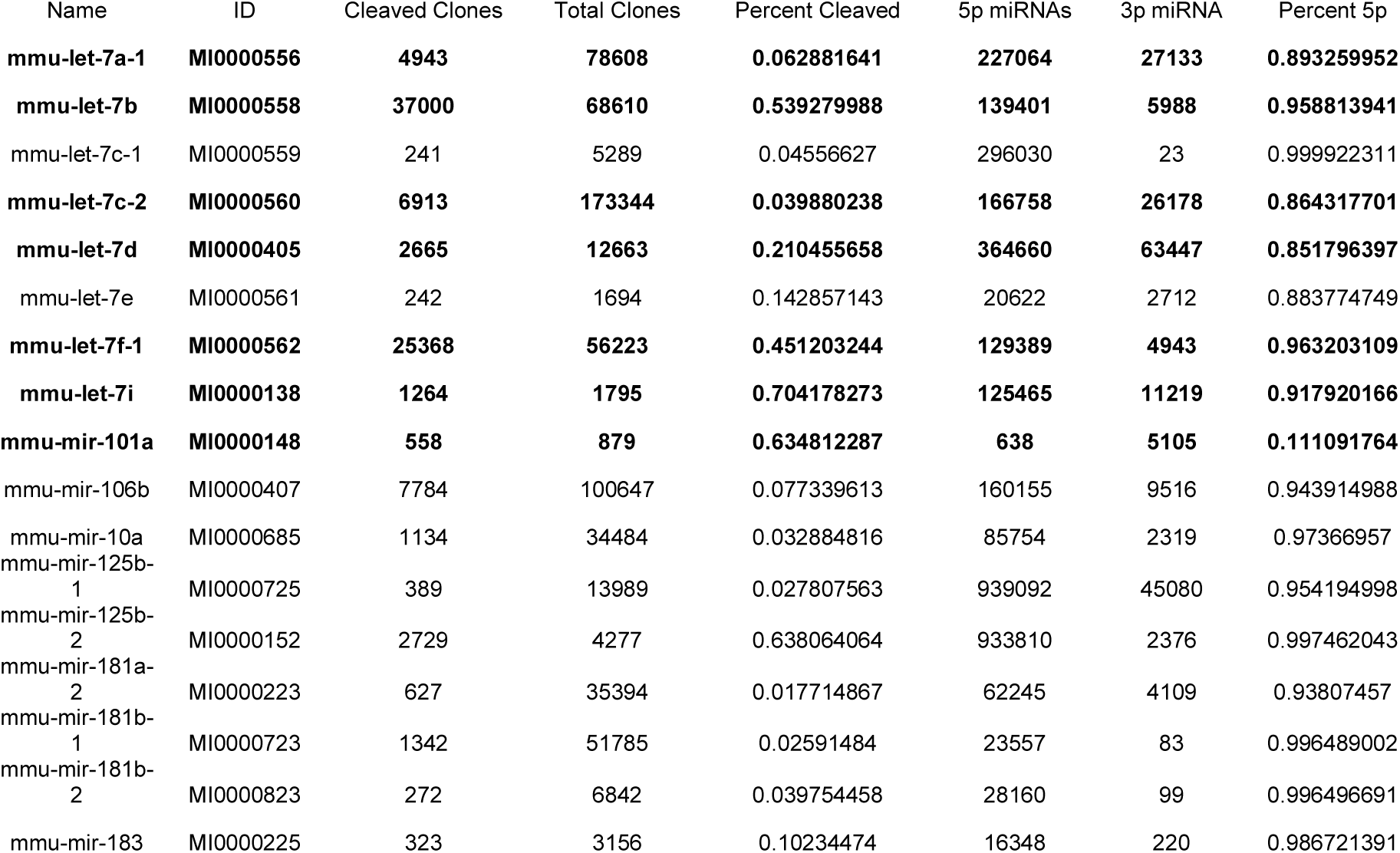
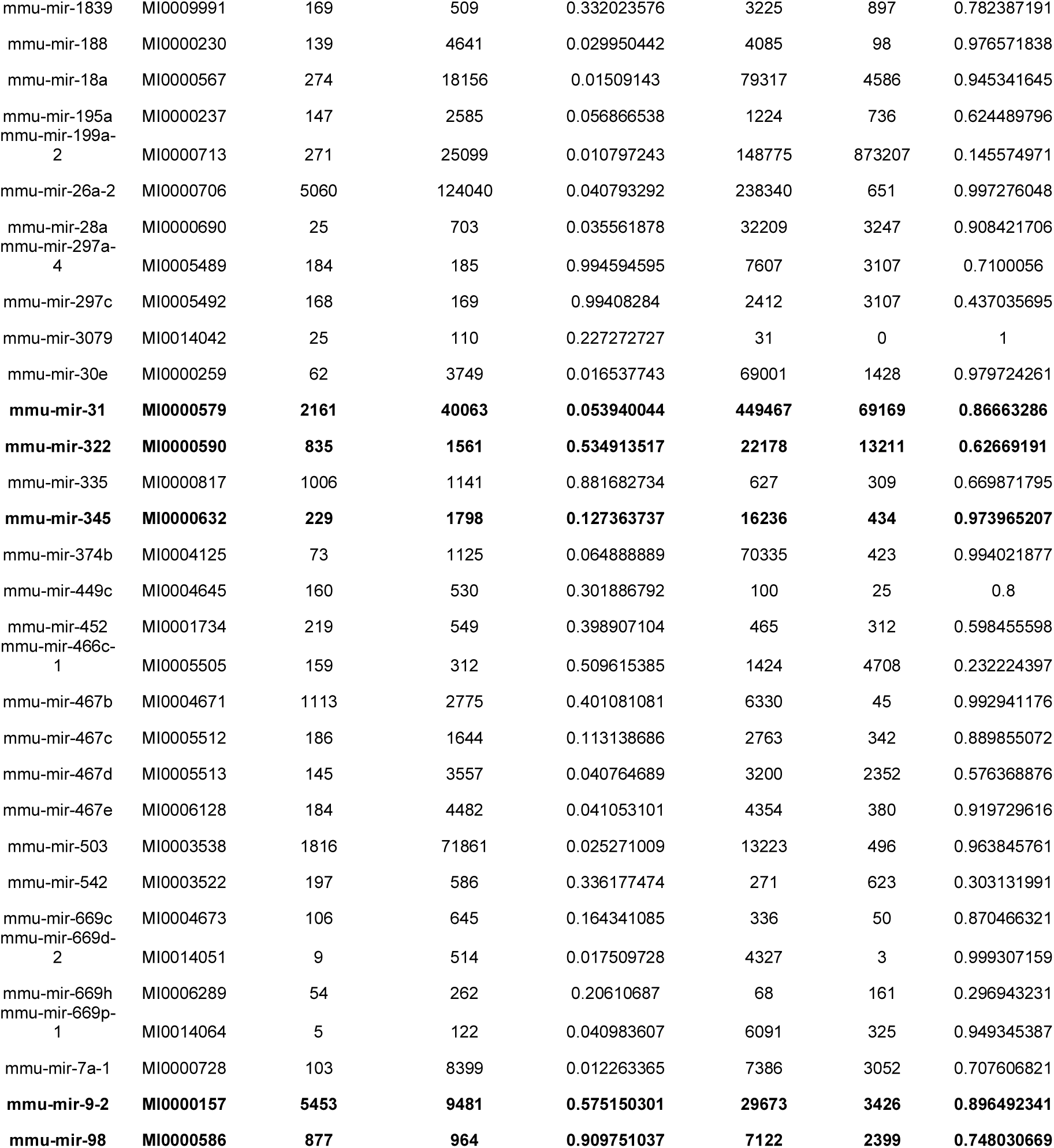
Known and putative ac-pre-miRNAs identified in MEFs

| Name | ID | Cleaved Clones | Total Clones | Percent Cleaved | 5p miRNAs | 3p miRNA | Percent 5p |
| --- | --- | --- | --- | --- | --- | --- | --- |
| **mmu-let-7a-1** | **MI0000556** | **4943** | **78608** | **0.062881641** | **227064** | **27133** | **0.893259952** |
| **mmu-let-7b** | **MI0000558** | **37000** | **68610** | **0.539279988** | **139401** | **5988** | **0.958813941** |
| mmu-let-7c-1 | MI0000559 | 241 | 5289 | 0.04556627 | 296030 | 23 | 0.999922311 |
| **mmu-let-7c-2** | **MI0000560** | **6913** | **173344** | **0.039880238** | **166758** | **26178** | **0.864317701** |
| **mmu-let-7d** | **MI0000405** | **2665** | **12663** | **0.210455658** | **364660** | **63447** | **0.851796397** |
| mmu-let-7e | MI0000561 | 242 | 1694 | 0.142857143 | 20622 | 2712 | 0.883774749 |
| **mmu-let-7f-1** | **MI0000562** | **25368** | **56223** | **0.451203244** | **129389** | **4943** | **0.963203109** |
| **mmu-let-7i** | **MI0000138** | **1264** | **1795** | **0.704178273** | **125465** | **11219** | **0.917920166** |
| **mmu-mir-101a** | **MI0000148** | **558** | **879** | **0.634812287** | **638** | **5105** | **0.111091764** |
| mmu-mir-106b | MI0000407 | 7784 | 100647 | 0.077339613 | 160155 | 9516 | 0.943914988 |
| mmu-mir-10a | MI0000685 | 1134 | 34484 | 0.032884816 | 85754 | 2319 | 0.97366957 |
| mmu-mir-125b-1 | MI0000725 | 389 | 13989 | 0.027807563 | 939092 | 45080 | 0.954194998 |
| mmu-mir-125b-2 | MI0000152 | 2729 | 4277 | 0.638064064 | 933810 | 2376 | 0.997462043 |
| mmu-mir-181a-2 | MI0000223 | 627 | 35394 | 0.017714867 | 62245 | 4109 | 0.93807457 |
| mmu-mir-181b-1 | MI0000723 | 1342 | 51785 | 0.02591484 | 23557 | 83 | 0.996489002 |
| mmu-mir-181b-2 | MI0000823 | 272 | 6842 | 0.039754458 | 28160 | 99 | 0.996496691 |
| mmu-mir-183 | MI0000225 | 323 | 3156 | 0.10234474 | 16348 | 220 | 0.986721391 |
| mmu-mir-1839 | MI0009991 | 169 | 509 | 0.332023576 | 3225 | 897 | 0.782387191 |
| mmu-mir-188 | MI0000230 | 139 | 4641 | 0.029950442 | 4085 | 98 | 0.976571838 |
| mmu-mir-18a | MI0000567 | 274 | 18156 | 0.01509143 | 79317 | 4586 | 0.945341645 |
| mmu-mir-195a | MI0000237 | 147 | 2585 | 0.056866538 | 1224 | 736 | 0.624489796 |
| mmu-mir-199a-2 | MI0000713 | 271 | 25099 | 0.010797243 | 148775 | 873207 | 0.145574971 |
| mmu-mir-26a-2 | MI0000706 | 5060 | 124040 | 0.040793292 | 238340 | 651 | 0.997276048 |
| mmu-mir-28a | MI0000690 | 25 | 703 | 0.035561878 | 32209 | 3247 | 0.908421706 |
| mmu-mir-297a-4 | MI0005489 | 184 | 185 | 0.994594595 | 7607 | 3107 | 0.7100056 |
| mmu-mir-297c | MI0005492 | 168 | 169 | 0.99408284 | 2412 | 3107 | 0.437035695 |
| mmu-mir-3079 | MI0014042 | 25 | 110 | 0.227272727 | 31 | 0 | 1 |
| mmu-mir-30e | MI0000259 | 62 | 3749 | 0.016537743 | 69001 | 1428 | 0.979724261 |
| **mmu-mir-31** | **MI0000579** | **2161** | **40063** | **0.053940044** | **449467** | **69169** | **0.86663286** |
| **mmu-mir-322** | **MI0000590** | **835** | **1561** | **0.534913517** | **22178** | **13211** | **0.62669191** |
| mmu-mir-335 | MI0000817 | 1006 | 1141 | 0.881682734 | 627 | 309 | 0.669871795 |
| **mmu-mir-345** | **MI0000632** | **229** | **1798** | **0.127363737** | **16236** | **434** | **0.973965207** |
| mmu-mir-374b | MI0004125 | 73 | 1125 | 0.064888889 | 70335 | 423 | 0.994021877 |
| mmu-mir-449c | MI0004645 | 160 | 530 | 0.301886792 | 100 | 25 | 0.8 |
| mmu-mir-452 | MI0001734 | 219 | 549 | 0.398907104 | 465 | 312 | 0.598455598 |
| mmu-mir-466c-1 | MI0005505 | 159 | 312 | 0.509615385 | 1424 | 4708 | 0.232224397 |
| mmu-mir-467b | MI0004671 | 1113 | 2775 | 0.401081081 | 6330 | 45 | 0.992941176 |
| mmu-mir-467c | MI0005512 | 186 | 1644 | 0.113138686 | 2763 | 342 | 0.889855072 |
| mmu-mir-467d | MI0005513 | 145 | 3557 | 0.040764689 | 3200 | 2352 | 0.576368876 |
| mmu-mir-467e | MI0006128 | 184 | 4482 | 0.041053101 | 4354 | 380 | 0.919729616 |
| mmu-mir-503 | MI0003538 | 1816 | 71861 | 0.025271009 | 13223 | 496 | 0.963845761 |
| mmu-mir-542 | MI0003522 | 197 | 586 | 0.336177474 | 271 | 623 | 0.303131991 |
| mmu-mir-669c | MI0004673 | 106 | 645 | 0.164341085 | 336 | 50 | 0.870466321 |
| mmu-mir-669d-2 | MI0014051 | 9 | 514 | 0.017509728 | 4327 | 3 | 0.999307159 |
| mmu-mir-669h | MI0006289 | 54 | 262 | 0.20610687 | 68 | 161 | 0.296943231 |
| mmu-mir-669p-1 | MI0014064 | 5 | 122 | 0.040983607 | 6091 | 325 | 0.949345387 |
| mmu-mir-7a-1 | MI0000728 | 103 | 8399 | 0.012263365 | 7386 | 3052 | 0.707606821 |
| **mmu-mir-9-2** | **MI0000157** | **5453** | **9481** | **0.575150301** | **29673** | **3426** | **0.896492341** |
| **mmu-mir-98** | **MI0000586** | **877** | **964** | **0.909751037** | **7122** | **2399** | **0.748030669** |

All previous examples of ac-pre-miRNAs have been shown to be cleaved within the 3p miRNA sequence and generate 5p miRNAs. To identify putative ac-pre-miRNAs that are processed with the opposite directionality, we performed a parallel search for pre-miRNA-seq reads that began 8-15 nucleotide from the 5’ end of the 5p miRNA. From this analysis, we found 3 candidates in humans and 5 candidates in mouse including hsa-miR-140 and mmu-miR-22 (Figs. 3C-D **and Table S3**). Importantly, these 5p cleaved ac-pre-miRNAs give rise predominantly to 3p miRNAs suggesting that cleavage in this region is not a degradation byproduct and is likely a competent mechanism for generating mature miRNAs (**Table S3**). These results reveal a greater collection of ac-pre-miRNAs than previously appreciated and uncover a likely novel class of 5p cleaved ac-pre-miRNAs.

### Relating pre-miRNA and mature miRNA abundance

Almost nothing is known about the relationship between pre-miRNA and mature miRNA abundance. In order to assess this, we first grouped miRNAs into their families and merged the mapped reads (**Table S4**). This was necessary to avoid artifacts, given that multiple distinct precursors give rise to identical, or nearly identical mature miRNAs. We further refined our analysis by only focusing on high confidence miRNAs, which left us with 158 human and 159 mouse miRNAs with reads mapping from pre-miRNA-seq and/or miRNA-seq libraries (**Table S5**). We analyzed the correlation between pre-miRNAs and mature miRNA levels expressed from these families (Figs. 4A-B). We found a positive but modest correlation between pre-miRNA and mature miRNA expression in both humans (Spearman correlation R = 0.53, p-value < 6.635e^−08^) and mouse (Spearman correlation R = 0.52, p-value < 2.047e^−07^). We also assessed the relationship between smRNA-seq and pre-miRNA-seq in HEK293T cells (Fig. S3A). We found a strikingly similar correlation to that of miRNA-seq (Spearman correlation; R = 0.53, p-value < 4.331e^−08^), which is explained by the high correlation of smRNA-seq to miRNA-seq (Spearman correlation; R = 0.90, p-value < 2.2e^−16^) (Fig. S3B). Together, these results reveal that in general pre-miRNA and mature miRNA levels have a modest positive correlation in both mouse and humans.

**Figure 4.**
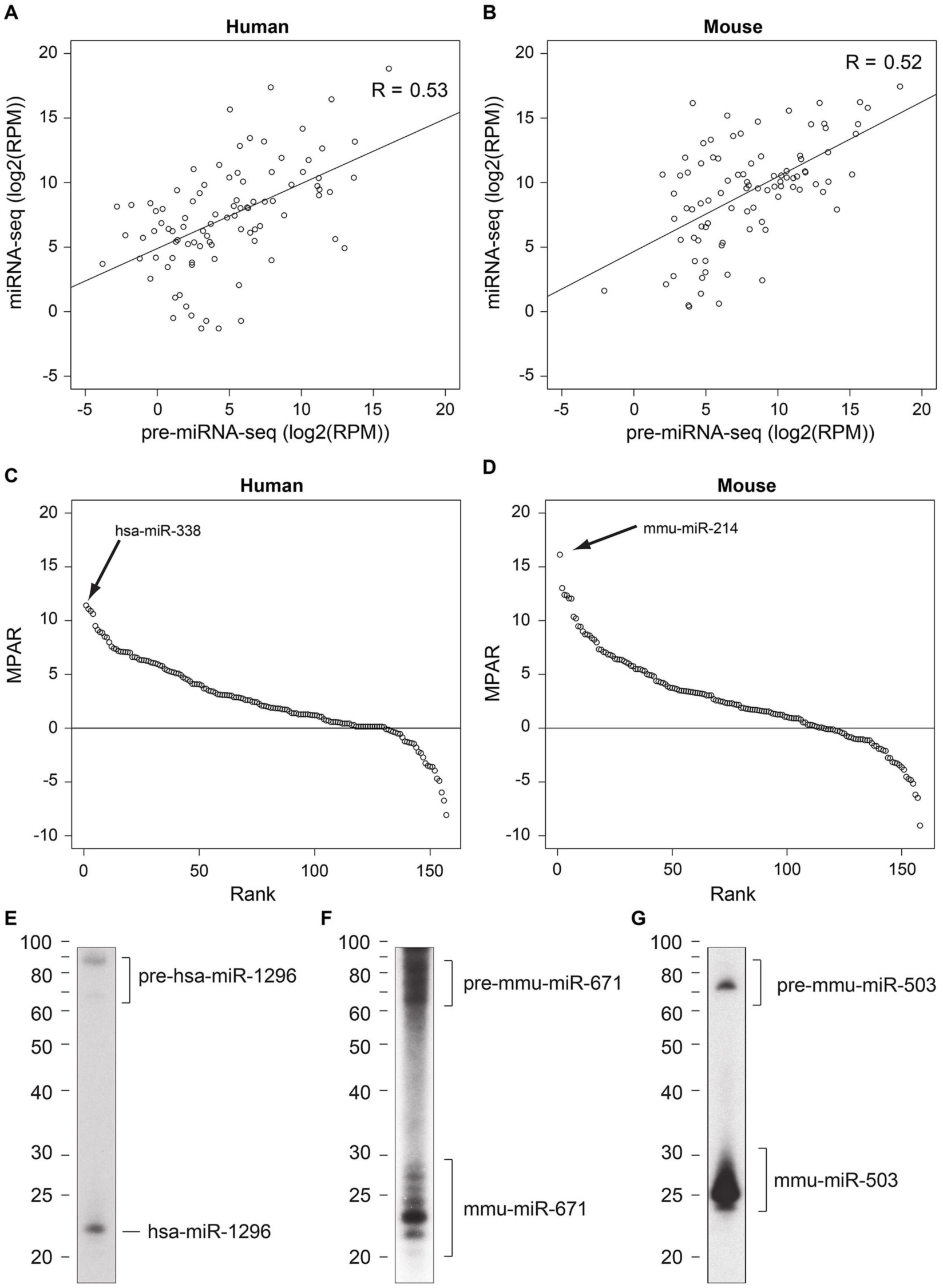
An index for miRNA precursor processing efficiency. A) Correlation of pre-miRNA-seq and miRNA-seq for high confidence human (A) and mouse (B) miRNA loci (human: R = 0.53, p-value < 6.635e^−08^, mouse: R = 0.52, p-value < 2.047e^−07^; Spearman correlation coefficient). C-D) Rank ordered list of mature to precursor abundance ratio (MPAR) scores for high confidence human (C) and mouse (D) miRNAs. E-G) Small RNA Northern blots of select miRNAs with low MPAR scores from human (HEK293T) or mouse (MEF) lysates.

### An index for miRNA precursor processing

We reasoned that the ratio of mature miRNAs to pre-miRNAs represents a reasonable estimate for *in vivo* miRNA processing efficiency. As some miRNAs had no detectable pre-miRNA reads or vice versa, we used a generalized log odds ratio of miRNA-seq to pre-miRNA-seq reads (**see METHODS**) to compute a mature to precursor abundance ratio (MPAR) for miRNAs expressed in human HEK293T and mouse MEF cells (**Tables S4 and S5**). As expected, the majority of high confidence miRNAs from human and mouse cells exhibited MPAR values > 0 (Figs. 4C-D and S3C), suggesting they are in general efficiently processed. Among the maximum scoring high confidence miRNAs in humans was hsa-miR-338, which had no detectable pre-miRNA-seq reads and 1,358 reads per million (RPM) in miRNA-seq libraries (MPAR = 11.4) (**Table S5**). In contrast, we found a number of miRNAs that had many more pre-miRNA-seq reads than mature miRNA-seq reads. To validate that our MPAR predictions are representative, we performed northern blotting on one human (hsa-miR-1296) and two mouse (mmu-miR-671 and mmu-miR-503) miRNAs with low MPAR scores (Figs. 4E-G). In each case, we observed bands corresponding to both the mature and precursor miRNA at similar levels. In comparison to the overall levels of pre-miRNA to mature miRNA (Fig. 1B) these miRNAs show a relative enrichment towards the pre-miRNA, indicating that they are inefficiently processed to mature miRNAs. Thus, examining the ratio of mature to pre-miRNAs allows us to approximate the efficiency of miRNA processing for hundreds of miRNAs in a single experiment.

### Distinct processing of two pre-miRNAs from Dgcr8 mRNA

The microprocessor complex (DGCR8/DROSHA) auto-regulates the expression of the *Dgcr8* transcript by binding and cleaving two hairpins near the 5’ end of its mRNA (52–54). We re-examined the abundance, modification status, and MPAR scores for the two annotated miRNAs in this region (Fig. S4). We determined that hsa-miR-3618, which is encoded in the 5’ untranslated region (UTR) of *Dgcr8*, is inefficiently processed into mature miRNAs (MPAR = −5.22), whereas hsa-miR-1306, which lies in the coding sequence (CDS), had a high MPAR score (MPAR = 3.09) (Figs. S4A-D **and Table S4**). Furthermore, we found that less than 10% of hsa-miR-3618 pre-miRNAs contained non-templated tails, whereas 100% of the hsa-miR-1306 clones were mono-uridylated, a signal that has previously been linked to efficient processing in some miRNAs (18). Thus, divergent processing of two miRNAs from the same primary transcript underscores the selectivity of DICER processing, and suggests that these two hairpins likely have evolved distinct functions in post-transcriptional regulation.

### Identification of AGO-associated stem-loops derived from mRNAs

Given this observation, we examined our dataset for novel pre-miRNAs embedded in other mRNAs. To do this, we took a conservative approach, using pre-miRNA-seq reads that failed to map to miRBase, and filtering them by mapping to small nuclear RNAs (snRNAs), snoRNAs, tRNAs, ribosomal RNAs (rRNAs), and repeat-masked sequences (Fig. 5A, **see Methods**). Approximately 1.2 and 0.37 million clones passed our stringent filtering steps and mapped to human and mouse mRNAs, respectively (**Table S1**). To identify pre-miRNA-like AGO-associated stem-loops derived from mRNAs, we used a CLIP-seq peak calling approach to identify significant (modified false discovery rate (mFDR) < 0.01) read clusters in mRNAs (35). We found a number of significant peaks that corresponded to nearly the full length of some highly expressed genes (e.g *ACTB*). Therefore, we further filtered significant clusters based on their size (< 200 nt), then chose the top clone from each cluster, folded it using RNAfold, and captured clusters that had a minimum free energy (MFE) < −0.3 kcal/mol/nt and a minimum of 15 base pairs (bp) in the longest hairpin (36).

**Figure 5.**
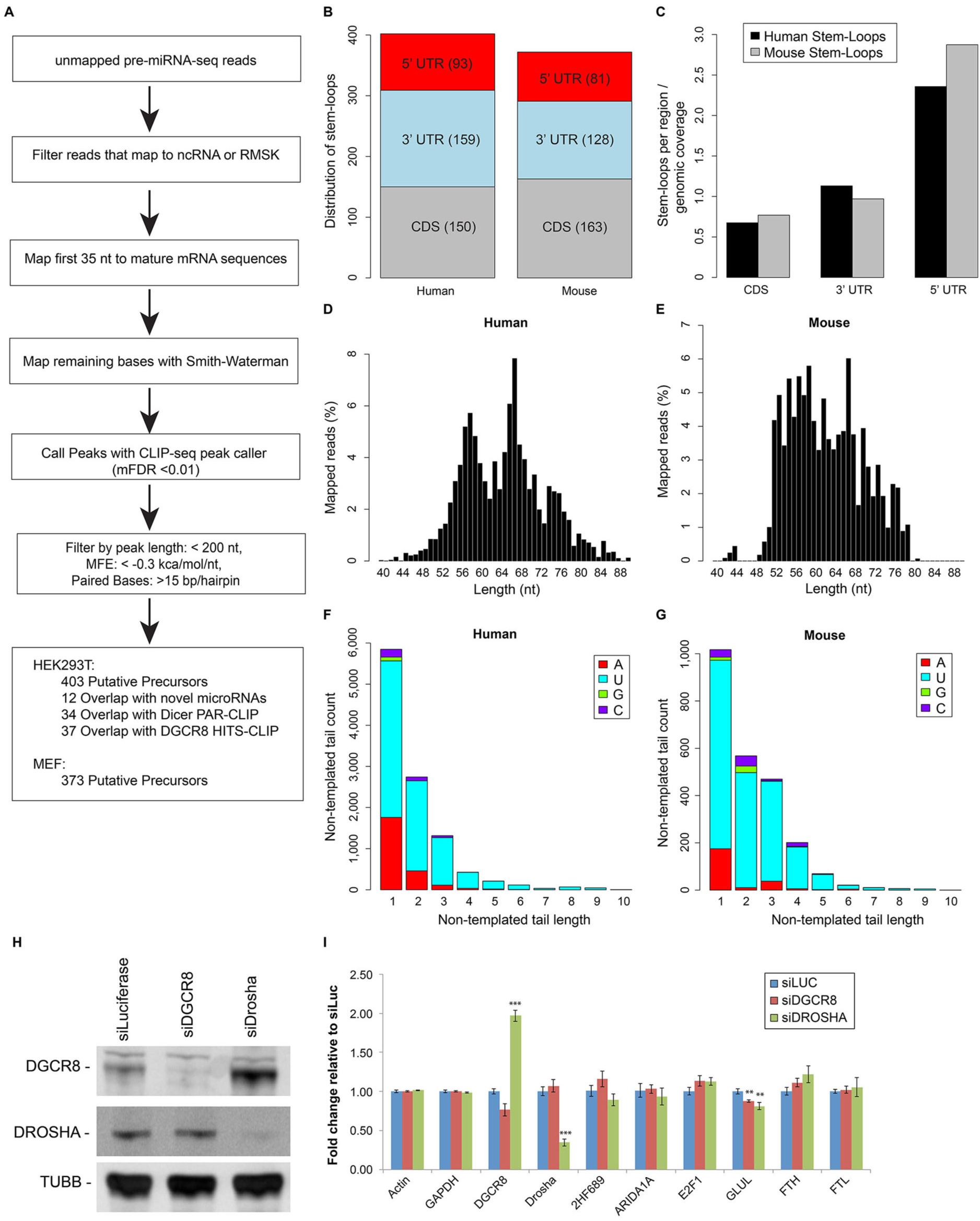
AGO-associated stem-loops derived from mRNAs. A) Bioinformatics pipeline for identification of AGO-associated stem-loops derived from mRNAs. Pre-miRNA-seq reads that did not map to miRBase were filtered on ncRNA and repeat-masker (RMSK), trimmed to 35 nt and mapped to spliced mRNA sequences. These alignments were then extended to find all matching bases. AGO-associated stem-loops derived from mRNAs were identified using a CLIP-seq peak caller (mFDR < 0.01) followed by length filtering (< 200 nt), MFE (< −0.3 kcal/mol/nt), and paired bases (> 15 bp/hairpin). B) Distribution of AGO-associated stem-loops derived from mRNAs. C) Enrichment of AGO-associated stem-loops derived from mRNAs within mRNA regions relative to their genomic coverage. D-E) Size distribution of pre-miRNA-seq reads from human (D) and mouse experiments (E) mapping to AGO-associated stem-loops derived from mRNAs. F-G) Non-templated 3’ end tail length and sequence content for pre-miRNA-seq reads mapping to human (F) and mouse (G) AGO-associated stem-loops derived from mRNAs. H) Western blot of *DGCR8*, *DROSHA*, and *TUBB* in HEK293T cells following siRNA knockdown. I) RT-qPCR analysis of mRNAs hosting AGO-associated stem-loops following knockdown of *DGCR8* or *DROSHA* in HEK293T cells. ** = p-value < 0.01, *** = p-value < 0.001; Students’ t-test.

This resulted in 403 and 373 AGO-associated stem-loops derived from human and mouse mRNAs, respectively (**Table S6**). We intersected our list of AGO-associated stem-loops derived from human mRNAs with recently identified miRNAs in humans from extensive smRNA-seq or from DICER photoactivatable-ribonucleoside-enhanced crosslinking and immunoprecipitation (PAR-CLIP) (26,55). In fact, 12 of our AGO-associated stem-loops were annotated as novel miRNAs in these lists supporting the ability of our approach to identify novel miRNAs (Fig. 5A). Furthermore, 34 and 37 AGO-associated stem-loops overlapped with DICER and DGCR8 binding sites respectively, revealing that a number of the AGO-associated stem-loops derived from mRNAs interact with other components of the canonical miRNA processing pathway (55,56).

We next examined the distribution of AGO-associated stem-loops derived from mRNAs across mRNA regions and found that they were equally present in all regions of mRNAs and similarly distributed between human and mouse (Fig. 5B). When we normalized the distribution of AGO-associated stem-loops derived from mRNAs by the relative genomic coverage of each mRNA region, we found that the CDS was underrepresented, whereas the 5’ UTR was enriched 2.5 fold for AGO-associated stem-loops in both organisms (Fig. 5C). We also examined the size distribution of reads mapping to AGO-associated stem-loops derived from mRNAs and found them to be similar in size in both mammals, between 55-75 nt in humans and 52-80 nt in mouse (Figs. 5D-E). This size range was slightly broader than pre-miRNA-seq reads that mapped to miRBase (Figs. 1E-F).

We also analyzed non-templated 3’ additions to pre-miRNA-seq reads mapped to AGO-associated stem-loops derived from human and mouse mRNAs (Figs. 5F-G **and Table S7**). We found that pre-miRNA-seq reads which mapped to these elements were enriched for uridylation events, and in fact, a higher percentage were oligo-tailed (13.0% in human and 24.3% in mouse), compared to reads mapping to miRBase (Figs. 2C and S2F). This result suggests that the AGO-associated stem-loops derived from mRNAs identified by our approach undergo similar modifications to known pre-miRNAs.

Finally, we overlaid *in vivo* RNA structure-probing data from mouse embryonic stem (ES) cells generated with *in vivo* click selective 2’-hydroxyl acylation and profiling (icSHAPE) onto our AGO-associated stem-loops derived from mouse mRNAs (Fig. S5). icSHAPE, chemically modifies the backbone of unpaired nucleotides, causing early termination of reverse transcriptase (57). Based on our RNAfold predictions, we examined the icSHAPE reactivity at paired or unpaired bases and found that icSHAPE reactivity was significantly higher at unpaired bases (Kruskal-Wallis one-way ANOVA; p-value < 8.18e^−133^) (Fig. S5A). This can been seen clearly when examining the RNAfold structure diagrams of AGO-associated stem-loops with icSHAPE reactivity overlaid (Figs. S5B-D). Together, our analyses provide numerous candidate pre-miRNA-like elements that are processed into 50-80 nt AGO-associated stem-loops from mRNAs in mammalian cells.

### AGO-associated stem-loop host mRNAs are generally not regulated by the microprocessor complex

To assess whether AGO-associated stem-loop containing mRNAs are regulated by the microprocessor complex, we performed siRNA-mediated knockdown of *DROSHA* and *DGCR8* in HEK293T cells and assessed changes in gene expression (Figs. 5H-J). We observed robust knockdown of both DROSHA and DGCR8 protein levels (Fig. 5H). Consistent with the role of DROSHA in regulating DGCR8 expression, we found that knockdown of *DROSHA* increased DGCR8 protein and RNA expression (Fig. 5H-I). To determine whether DGCR8 and/or DROSHA participate in the regulation of AGO-associated stem-loop containing mRNAs, we performed mRNA-seq and RT-qPCR on knockdown RNA samples. To our surprise, the majority of mRNAs hosting AGO-associated stem-loops were unaffected by knockdown of components of the microprocessor. (**Table S7 and** Fig. 5I). In fact, the only gene tested by RT-qPCR that was significantly affected was *GLUL,* but knockdown of either DGCR8 or DROSHA decreased the level of *GLUL* expression, in contrast to the expected role of the microprocessor. These data suggest that cleavage of AGO-associated stem-loops does not have a significant impact on the expression of the host mRNA because cleavage events are rare, or because this process is independent of the microprocessor, with neither model being mutually exclusive.

### Most AGO-associated stem-loops derived from mRNAs do not produce AGO-bound smRNAs

We next calculated the miRNA-seq and smRNA-seq coverage at AGO-associated stem-loops derived from mRNAs. We found that miRNA-seq and smRNA-seq reads mapping to AGO-associated stem-loops derived from mRNAs were of a similar size as miRBase miRNAs and had non-templated additions to their 3’ ends (Figs. 6A-B and S6A-D). However, when we calculated the MPAR scores for AGO-associated stem-loops derived from mRNAs, we found that they had overall very low scores, indicating that they are poor substrates for producing mature AGO-bound smRNAs (Figs. 6C and S6E-F **and Table S8**). This is in agreement with the finding that these elements are more commonly oligo-tailed compared to canonical pre-miRNAs. In fact, we only found a total of 171 miRNA-seq reads mapping to AGO-associated stem-loops derived from human mRNAs and the majority of these reads mapped to the recently identified miRNAs embedded within the mRNAs; *GNAS, GLUL*, and *E2F1* (26,55) (example of E2F1 in Figs. 6D-E). Additionally, we found a total of 2,240 smRNA-seq reads mapping to AGO-associated mRNA stem-loops derived from human mRNAs, which in part reflects the higher sequencing depth of these libraries. Some of stem-loops with the most smRNA-seq reads, corresponded to previously identified novel miRNAs, including *BRD2*, *GLUL* and *E2F1* (**example in** Fig. 6F). However, we also noticed numerous reads mapping to mRNA stem-loops that did not have binding evidence from miRNA-seq, including *FTH1*, *FTL*, *SOX4*, and *KMT2C*.

**Figure 6.**
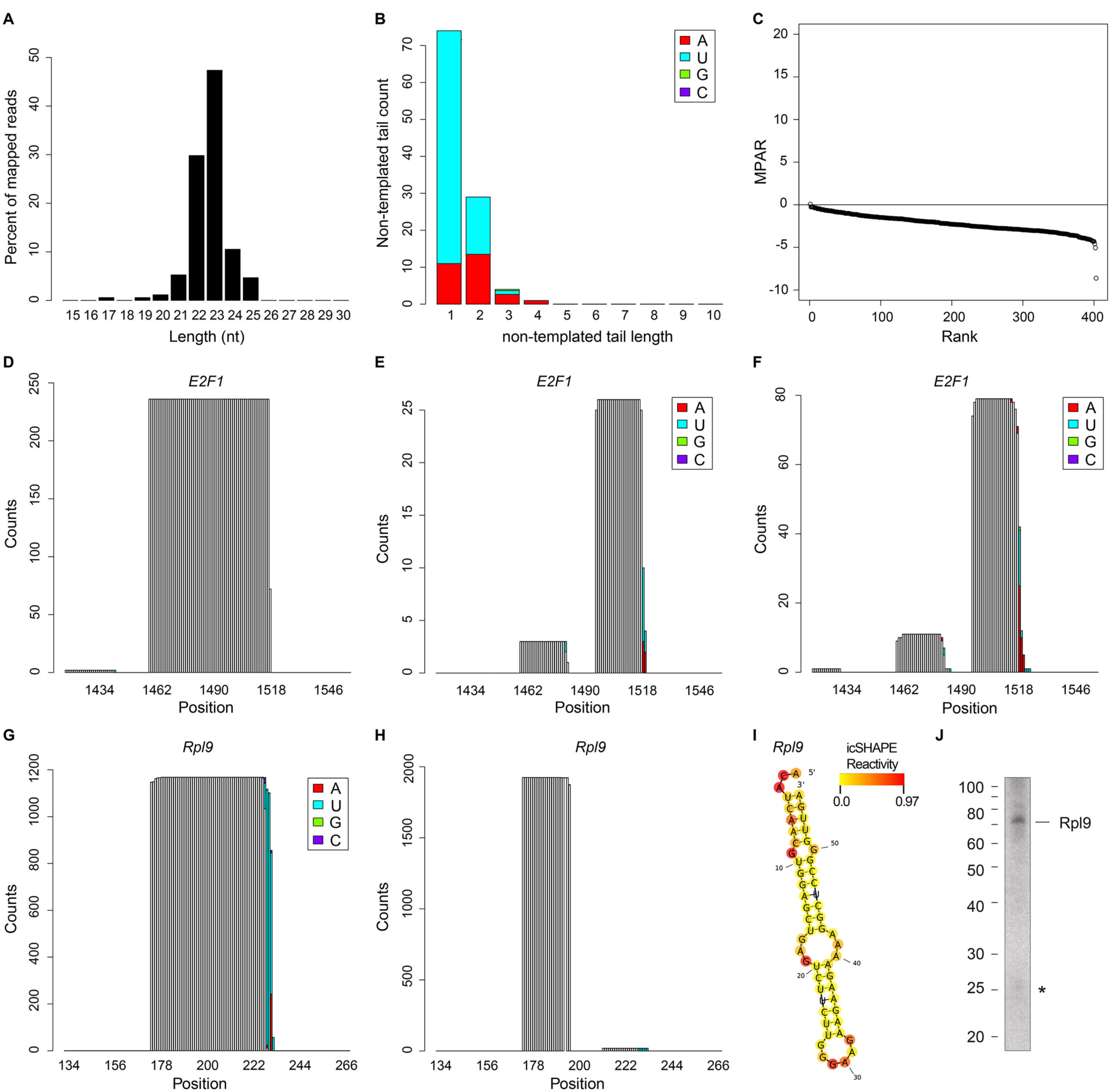
Few AGO-associated stem-loops derived from mRNAs produce smRNAs. A) Size distribution of miRNA-seq reads mapping to human AGO-associated stem-loops derived from mRNAs. B) Non-templated 3’ end tail length and sequence content for miRNA-seq reads mapping to human AGO-associated stem-loops derived from mRNAs. C) Rank order of MPAR score for AGO-associated stem-loops derived from human mRNAs. D-F) Coverage plot of pre-miRNA-seq (D), miRNA-seq (E), and smRNA-seq (F) reads mapping to a human AGO-associated stem-loop in the 3’ UTR of *E2F1*. White bars indicate templated nucleotides, colored bars indicate non-templated additions. G-H) Coverage plot of pre-miRNA-seq (G) and miRNA-seq (H) reads mapping to a mouse AGO-associated stem-loop in the CDS of *Rpl9*. White bars indicate templated nucleotides, colored bars indicate non-templated additions. I) RNAfold predicted structure and icSHAPE reactivity for mouse AGO-associated stem-loop in the CDS of *Rpl9*. J) Small RNA Northern blot of mouse AGO-associated stem-loop in the CDS of *Rpl9* from mouse (MEF) cells. * indicates band at the expected size of miRNA-like product.

For AGO-associated stem-loops derived from mouse mRNAs we mapped 2,698 miRNA-seq reads, however 71% of these mapped to a single locus in the CDS of *Rpl9* (Figs. 6G-I). Northern blotting with a probe complementary to this sequence revealed a strong signal at ~70 nt, corresponding to the stem-loop, and a weak signal at 25 nt, corresponding to the expected size of the novel miRNA. (Fig. 6J). Therefore, we provide evidence that *Rpl9* is processed into a stem-loop like structure and further processed, albeit with low efficiency, into a miRNA-like smRNA. Other AGO-bound smRNA producing stem-loops were regions of the *Cyr61* 5’ UTR, *Asf1b* 5’ UTR, and *Klf9* 3’ UTR (Fig. S7). Collectively, these data demonstrate that pre-miRNA-seq uncovers AGO-associated mRNA stem-loops, some of which may represent novel miRNAs, but most of which are poorly processed into AGO-bound smRNAs in the cell lines investigated in this study.

### Iron response elements are processed into AGO-associated stem-loops

The top AGO-associated stem-loop candidate in humans is derived from the 5’ UTR of Ferritin heavy chain (*FTH1*) mRNA (Fig. 7A). We identified over 5,000 pre-miRNA-seq reads in this region, which had a strong predicted hairpin structure, clearly defined ends, and significant mono-tailing on the 3’ end. Intriguingly, this region of the *FTH1* transcripts corresponds precisely to the iron responsive element (IRE), which is a well-studied structural element that regulates translation in an iron-dependent fashion (58). We scanned our list of AGO-associated stem-loops for other IRE containing genes, and found Ferritin light chain (*FTL*) mRNA was also producing AGO-associated stem-loops from its IRE region (Fig. 7E). Furthermore, we identified AGO-associated stem-loops supported by icSHAPE data from the mouse homologs of both of these genes, *Fth1* and *Ftl* (Figs. S8A-E). Finally, we confirmed the presence of the *FTH1* pre-miRNA-like structure by northern blotting (Fig. 7D). Therefore, processing of IREs from ferritin genes into AGO-associated stem-loops is conserved in mammals.

**Figure 7.**
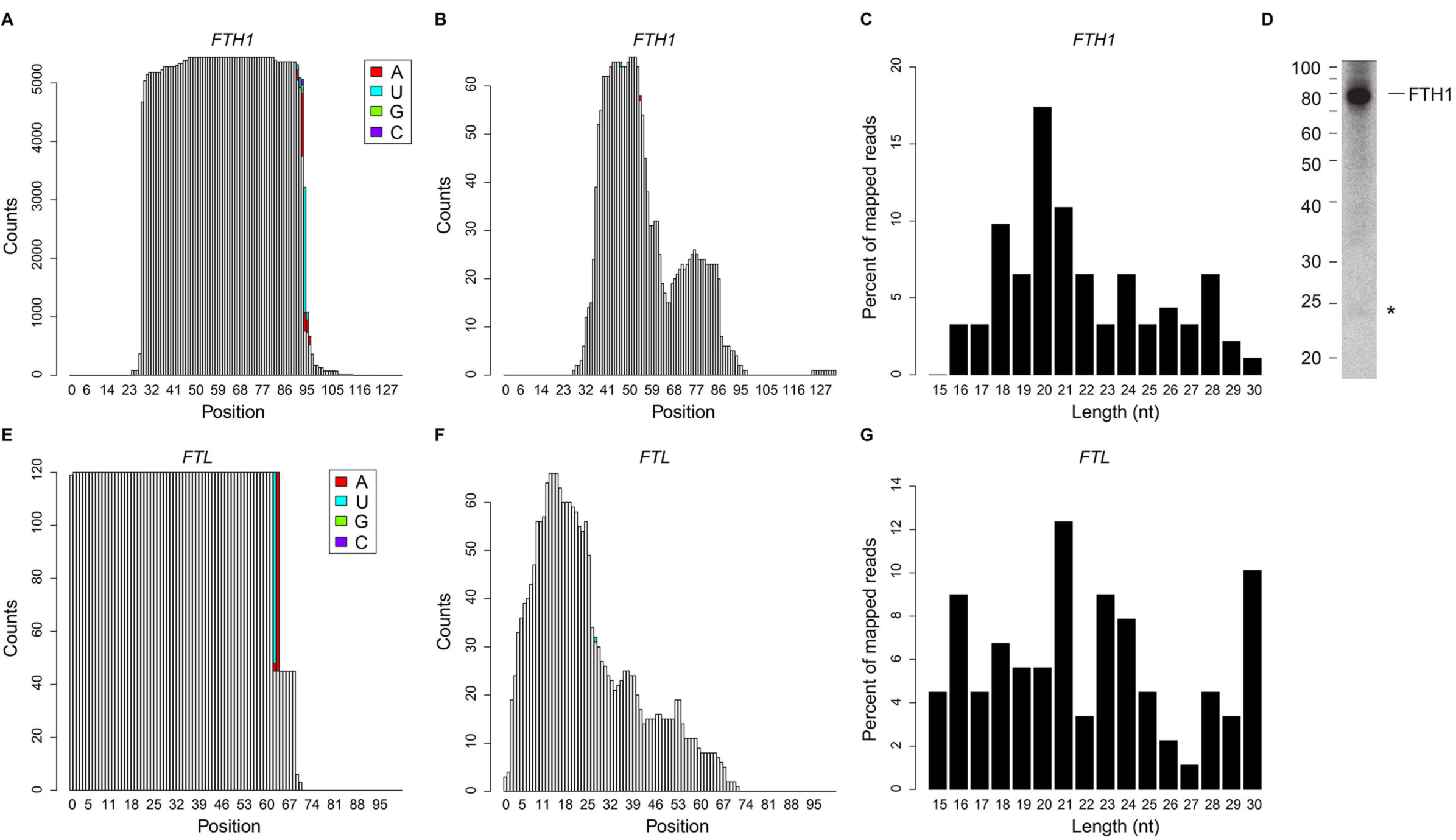
The IREs of human *FTH1* and *FTL* are processed into AGO-associated stem-loops. A-B) Coverage plot of pre-miRNA-seq (A) and smRNA-seq (B) reads mapping to a human AGO-associated stem loop in the 5’ UTR of the *FTH1* gene. White bars indicate templated nucleotides, colored bars indicate non-templated additions. C) Size distribution of smRNA-seq reads mapping to the human AGO-associated stem-loop in the 5’ UTR of *FTH1*. D) Small RNA Northern blot of human AGO-associated stem-loop in the 5’UTR of *FTH1* from HEK293T cells. * indicates band at the expected size of miRNA-like product. E-G) Coverage plot of pre-miRNA-seq (E) and smRNA-seq (F) reads mapping to a human AGO-associated stem loop in the 5’ UTR of the *FTL* gene. White bars indicate templated nucleotides, colored bars indicate non-templated additions. G) Size distribution of smRNA-seq reads mapping to the human AGO-associated stem-loop in the 5’ UTR of *FTL*.

To examine whether IRE-processed hairpins produce functional small RNAs, we examined miRNA-seq and smRNA-seq data from these regions. We did not find any AGO-interacting small RNAs from human *FTH1* or *FTL*. Nevertheless, we did observe cellular smRNAs from these regions in our smRNA-seq data (Figs. 7B,F). However, they were heterogeneous in size and only loosely reflective of DICER processing (Figs. 7C,G). Furthermore, smRNAs corresponding to the FTH1 were detectable by northern blotting but with much lower levels than the pre-miRNA-like structure. (Fig. 7D). Therefore, these smRNAs appear to be rare and likely the consequence of subsequent degradation of the processed stem-loops that are not loaded into AGO to make a functional RISC complex. In mouse, miRNA-seq only demonstrated a small number of clones originating from the *Fth* pre-miRNA (Figs. S8D), further corroborating our results from human cells. Intriguingly, siRNA-mediated knockdown of *DROSHA* or *DGCR8* in HEK23T cells had no effect on *FTH1* or *FTL* mRNA expression levels (Fig. 5I). This suggests that processing of IREs into stem-loops is DROSHA-independent and may work through a different endonuclease. In total, these results demonstrate that mammalian IREs are processed into AGO-bound stem-loops through a microprocessor-independent mechanism, and that these stem-loops are unlikely to be substrates for DICER. Whether cleaved IRE stem-loops are functional molecules remains to be determined.

## DISCUSSION

Here, we describe the application and further development of a methodology to enrich for and sequence AGO-associated pre-miRNAs in both human and mouse cells. This biochemical approach combined with custom bioinformatics pipelines successfully enriches for and maps pre-miRNAs in mammalian genomes (Fig. 1). Using this approach, we detected 367 pre-miRNAs in human and 267 in mouse cell lines, with ~ 28 and ~ 17 million raw sequencing reads in each experiment, respectively (**Tables S1 and S2**). This gave us specific insights into the exact sequence and abundance of pre-miRNAs and miRNAs expressed in cells of two different mammalian organisms. We generated profiles to visualize coverage, trimming, and non-templated tailing at each annotated miRNA expressed in either cell type, which is available for download at http://gregorylab.bio.upenn.edu/AGO_IP_Seq/.

Using these unique datasets, we uncovered widespread trimming and non-templated tailing in both pre- and mature miRNAs from human and mouse cells (Figs. 2, S1C-D and S2D-G). We also identified known and putative ac-pre-miRNAs (Figs. 3A-B and Tables 1 **and S3**). The large number of ac-pre-miRNAs identified suggests that DICER-independent pre-miRNA processing may be a more commonly used mechanism than previously appreciated (13,16,19). Furthermore, we identified putative ac-pre-miRNAs that cleave in the 5p arm of the pre-miRNA (**Table S3 and** Figs 3C-D), and thus are processed in the opposite direction of the currently known members of this pre-miRNA class. This potentially novel pre-miRNA processing mechanism would require an alternative maturation process, with processive exonucleolytic nucleotide removal occurring step-wise from the 5’ end.

Given the unique nature of our datasets, we were able to make the first comprehensive analysis of the relationship between pre-miRNA and mature miRNA abundance in an unbiased fashion (Figs. 4A-B and S3A). Remarkably, we found very consistent relationships between mature miRNA and pre-miRNA abundance in human and mouse cells. We determined a mature to precursor abundance ratio (MPAR), allowing us to investigate productive and unproductive miRNA maturation (Figs. 4C-D and S3C). We uncovered some pre-miRNAs that make surprisingly few mature species, despite abundant precursors. We also found that two miRNAs in *DGCR8* mRNA are processed with highly divergent efficiencies, suggesting distinct functionalities (Fig. S4).

From a further examination of pre-miRNA-seq reads that did not map to miRBase, we identified AGO-associated stem-loops derived from mRNAs (Fig. 5A). We found that these were enriched in the 5’ UTR of these transcripts, which suggests they may play a similar role to miR-3618 in the 5’ UTR of human *DGCR8* (Figs. 5B-C). Furthermore, we found that these stem-loops had a broader size distribution than known miRNAs and were oligo-uridylated, suggesting they are processed by TUTases (Figs. 5D-G). RNA structure prediction algorithms and *in vivo* structure probing data from mouse ES cells provide strong evidence that these regions of mRNAs form stem-loops (Fig. S5). Surprisingly, AGO-bound stem-loop containing mRNAs were not impacted following siRNA-mediated knockdown of components of the microprocessor complex (Fig. 5I). Our results strongly support the presence of AGO-associated stem-loops from mRNAs, however their biogenesis pathway remains undetermined. Furthermore, we found limited evidence for further processing into mature miRNA-like smRNAs, which may be cell-type or contextdependent and thus should be the target of future investigations.

Interestingly, we found very few mature AGO-bound sequences coming from these regions and overall low MPAR scores (Figs. 6A-C and S6). In human cells, the vast majority of AGO-bound smRNAs from these regions can be explained by recently identified novel miRNAs (26,55). In mouse cells, we uncovered a region in the *Rpl9* CDS and a few other putative miRNAs that account for most of the AGO-bound smRNAs from these regions (Figs. 6G-I and S7). These findings indicate that our methodology uncovers novel miRNAs (Fig. S7), but the function of most of these stem-loops, which produce few AGO-bound miRNAs, remains a mystery.

We also found that the IRE elements of human and mouse ferritin mRNAs are processed into pre-miRNA-like molecules, but rarely into mature AGO-bound smRNA species (Figs. 7 and S8). Further processing of theses hairpins may only occur in specific cell types (i.e. blood cells) under stress conditions. Moreover, knockdown of *DROSHA* or *DGCR8* had no effect on the expression of IRE host genes (Fig. 5I). Then, what is the function of these pre-miRNA-like molecules? IRE hairpins may represent stable remnants of normal degradation of IRE containing mRNAs. However, given the conservation of AGO-associated stem-loops from IREs in both human and mouse cell, this seems unlikely. Alternatively, they may be processed from ferritin host genes by endonucleases other than the DGCR8/DROSHA microprocessor. This would also be consistent with the inability of these pre-miRNA-like molecules to serve as substrates for DICER processing. However, these stem-loops could serve alternative roles, such as acting as an RNA-binding protein sink for IRE-binding proteins or other factors. Undoubtedly, the biogenesis and biological relevance of the IRE and other processed stem-loops will be the subject of further investigation.

## DATA AVAILABILITY

All sequencing data generated in this study has been deposited in GEO under the accession number GSE71710 (available at http://www.ncbi.nlm.nih.gov/geo/). RNAfold diagrams and coverage plots are available for download at http://gregorylab.bio.upenn.edu/AGO_IP_Seq/. MEF pre-miRNA-seq and miRNA-seq data were downloaded from European Nucleotide Archive (http://www.ebi.ac.uk/ena/) under the accession number PRJEB6756. HEK293T smRNA-seq data was obtained from GSE66224.

## ACKNOWLEDGMENTS

The authors thank members of the Gregory and Mourelatos laboratories for helpful discussions.

---

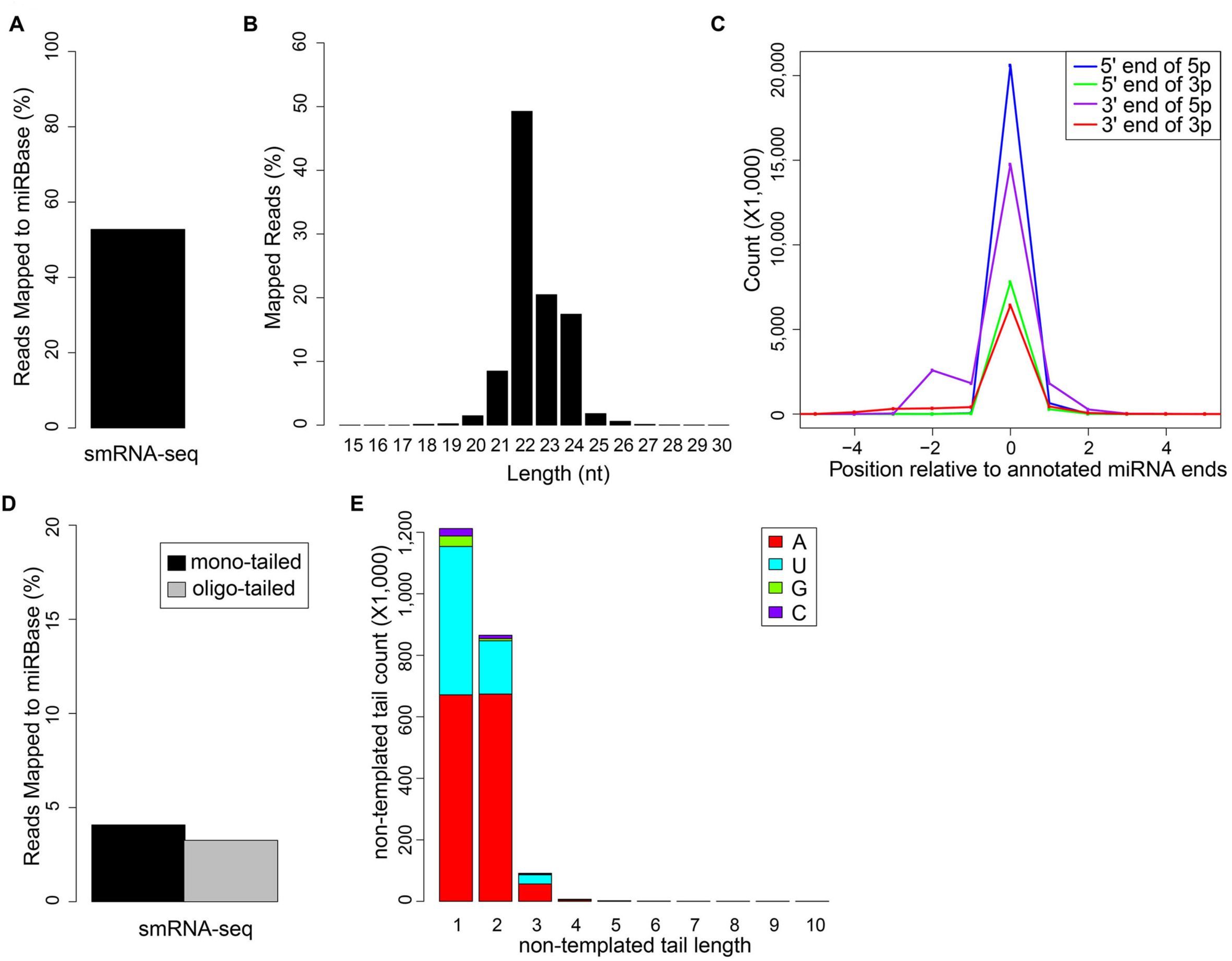

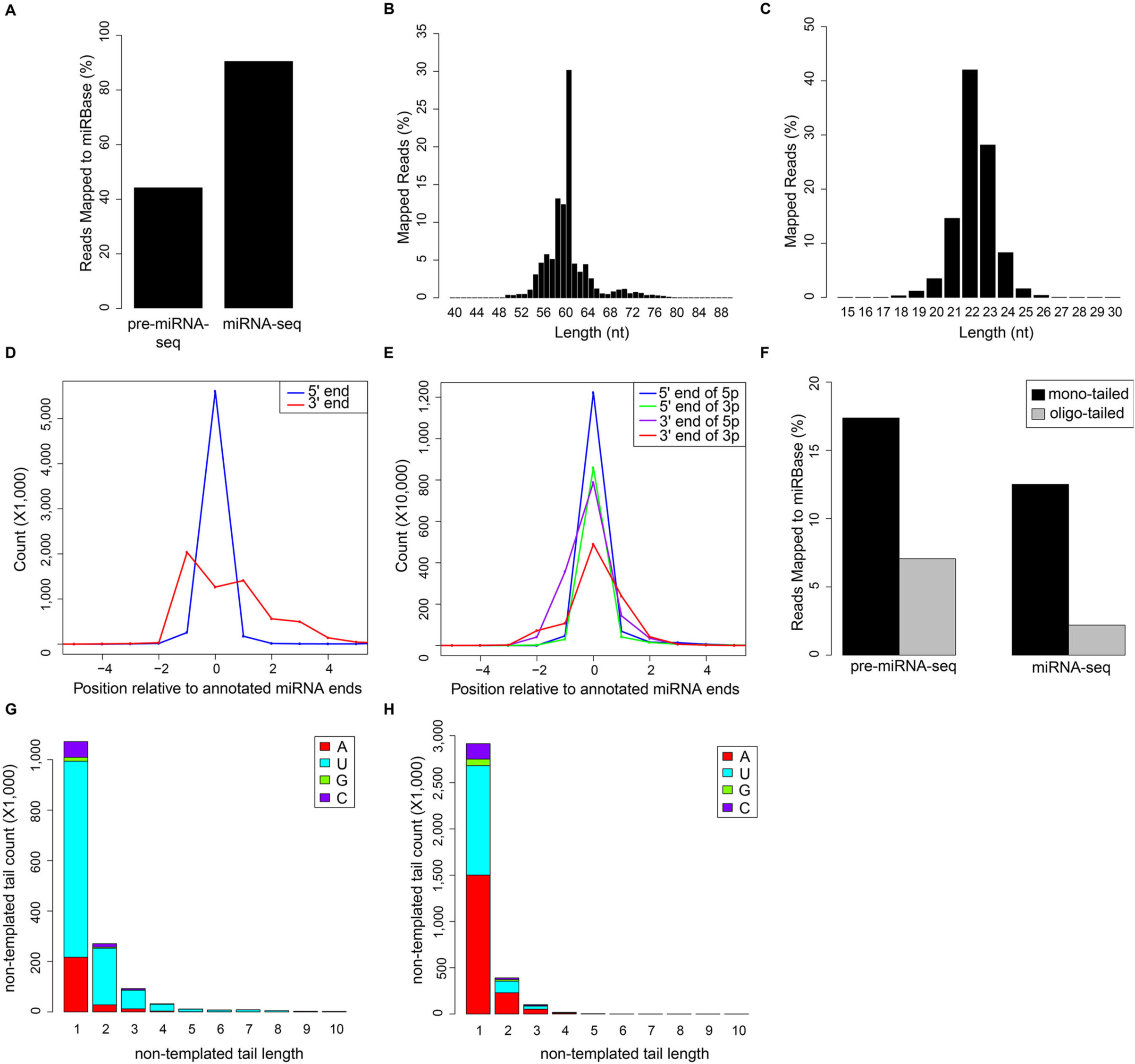

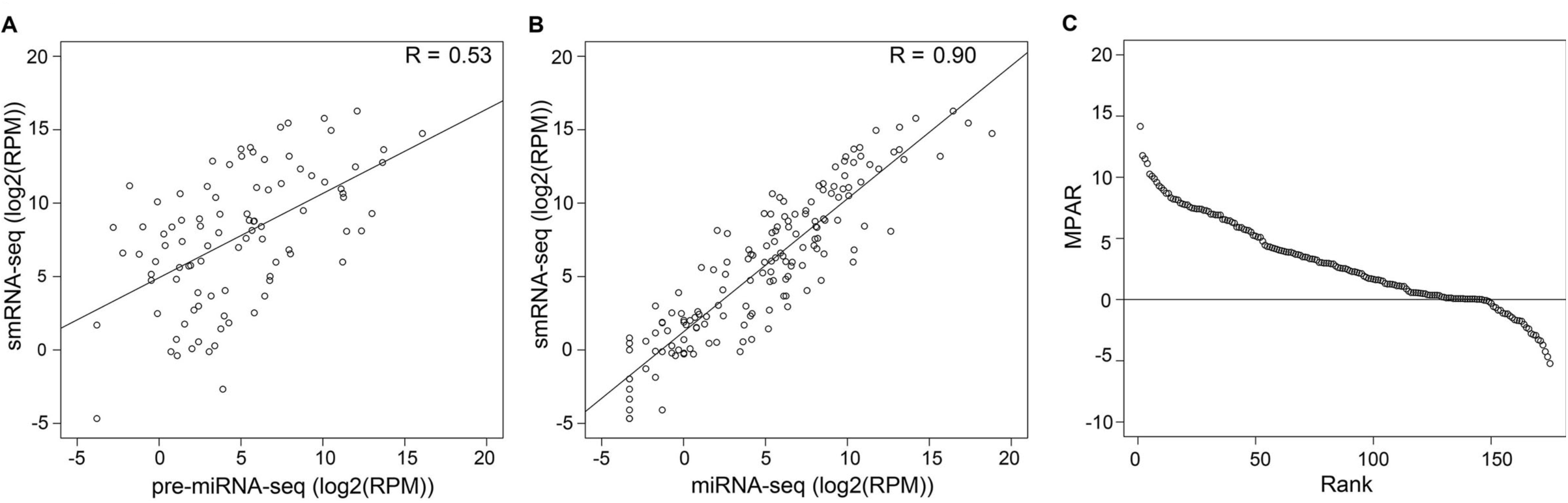

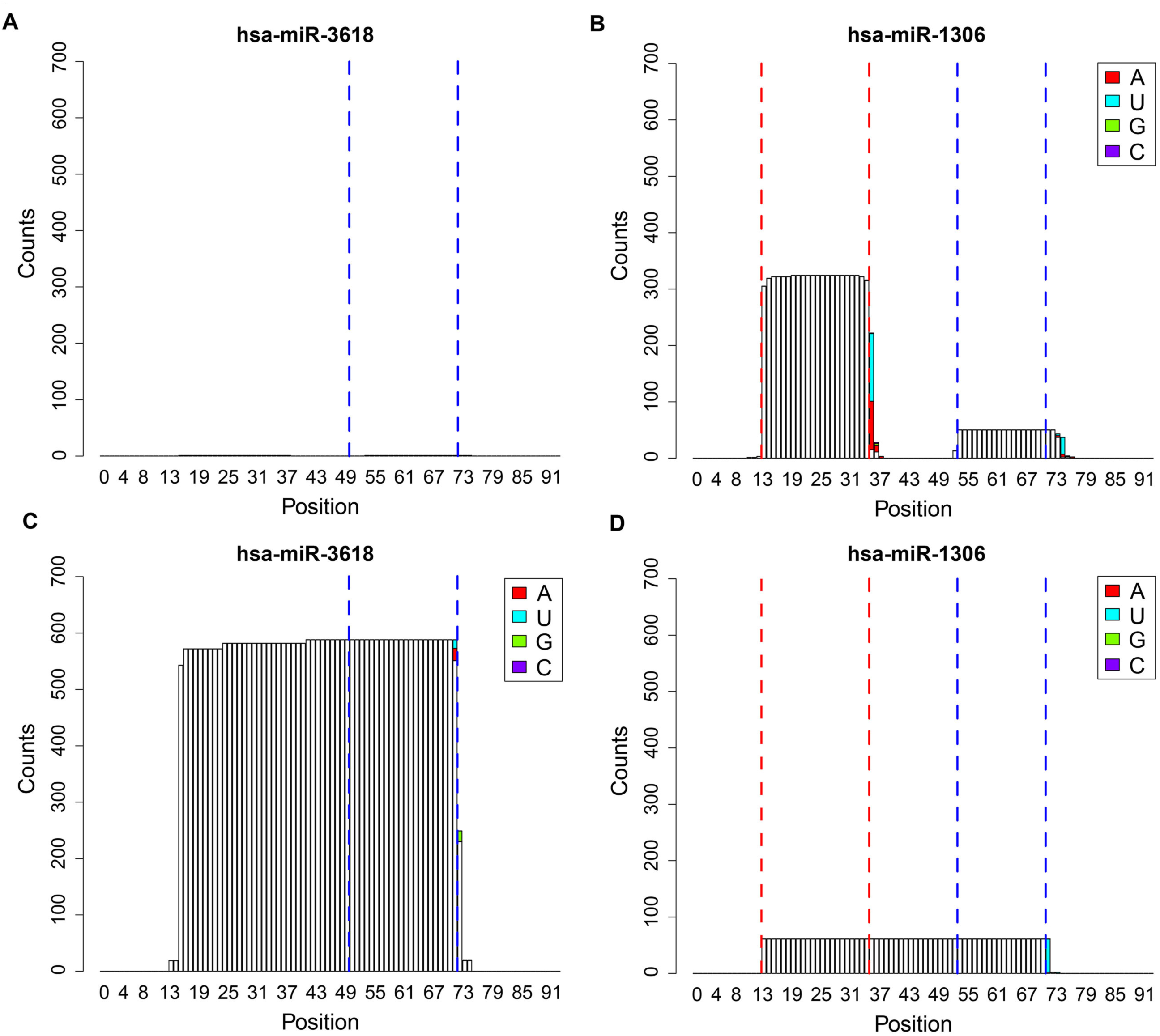

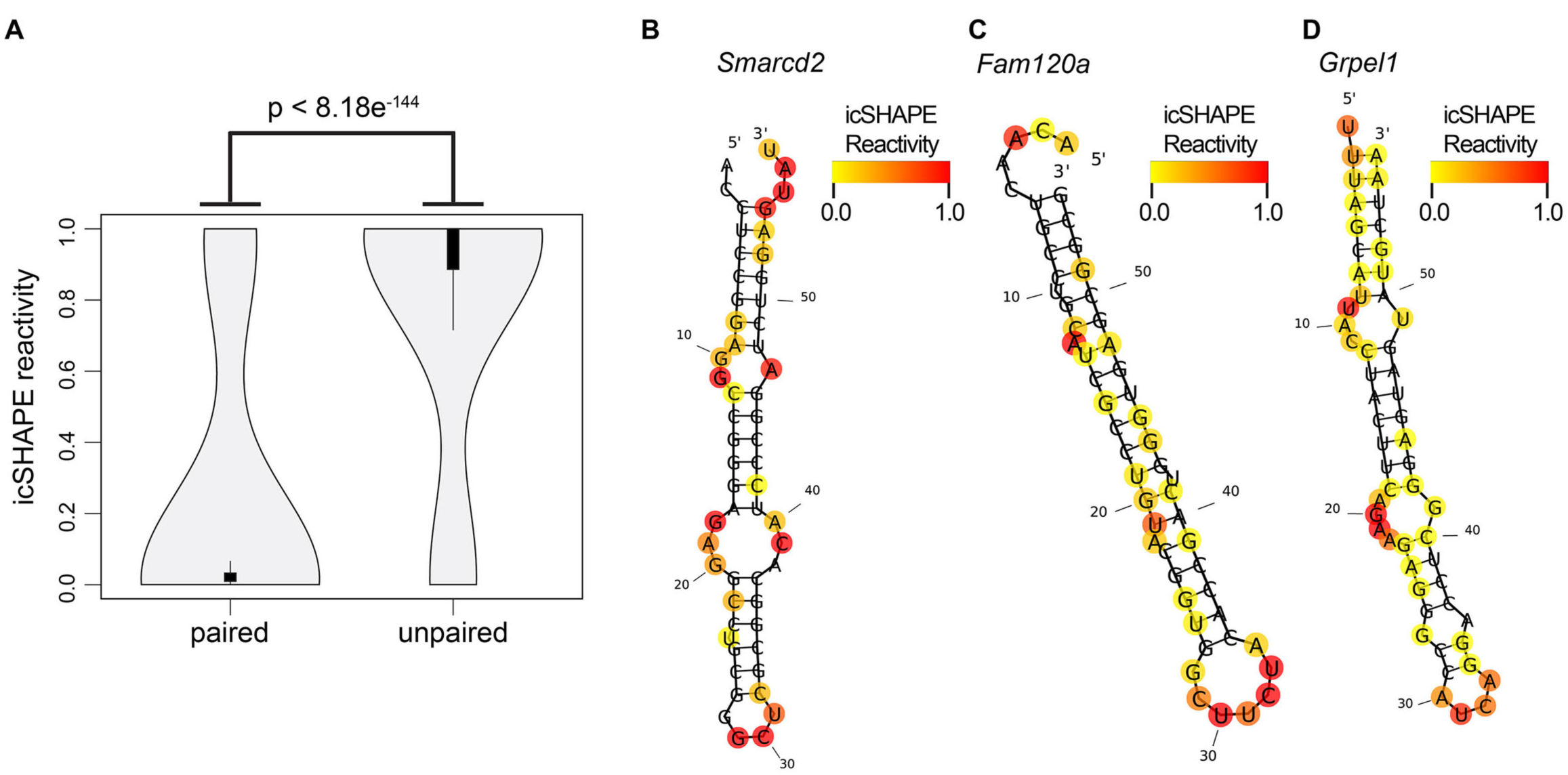

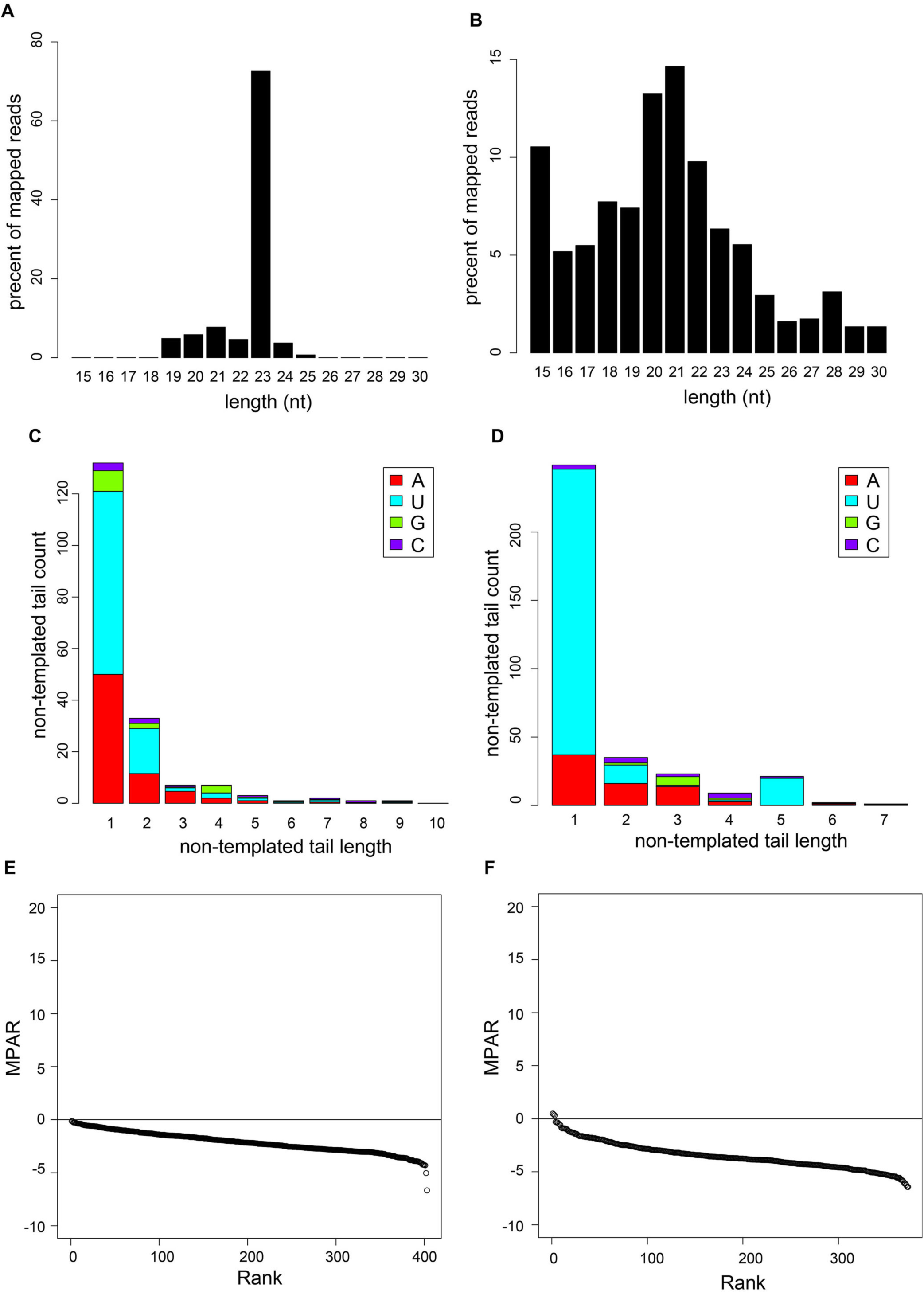

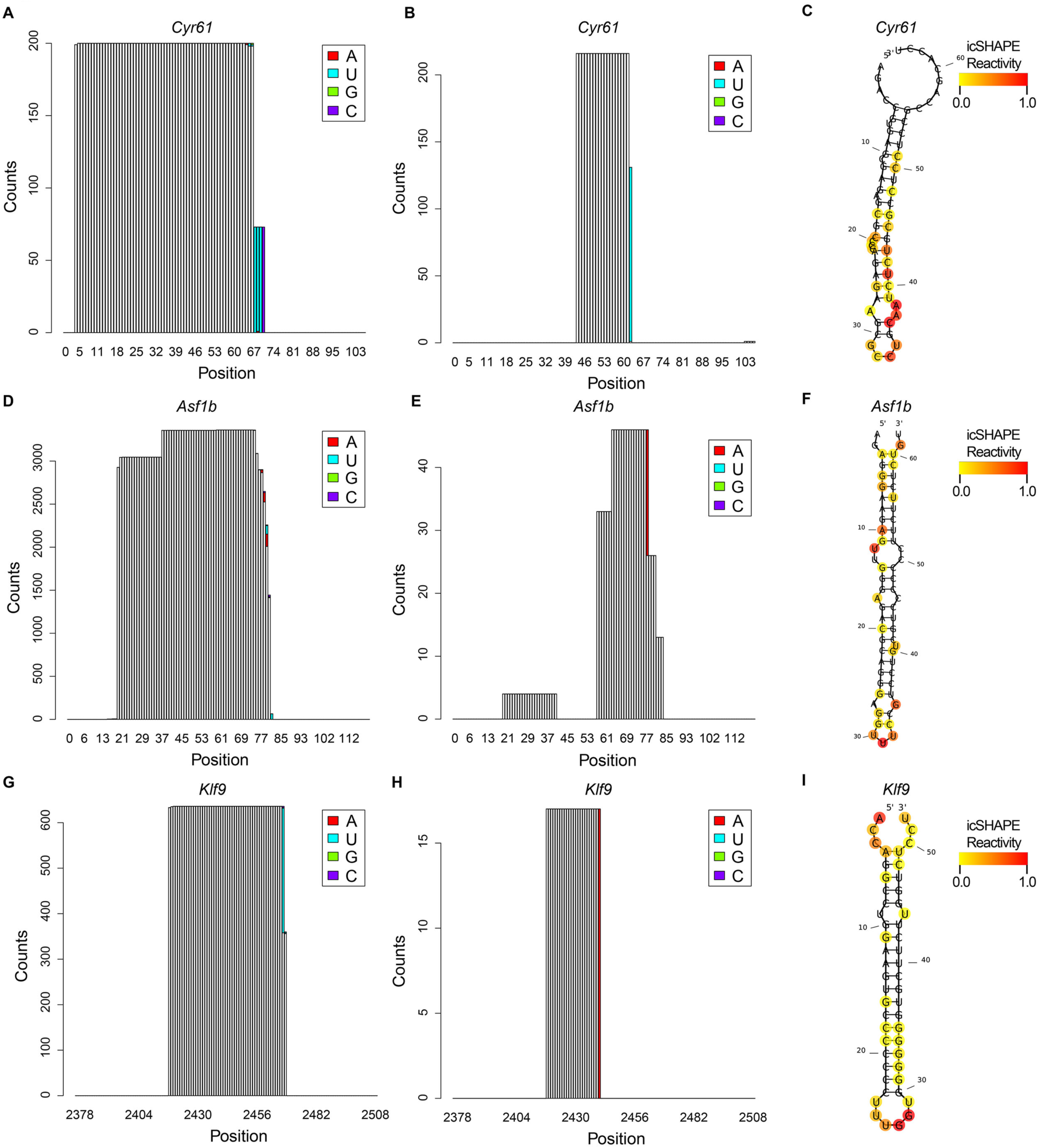

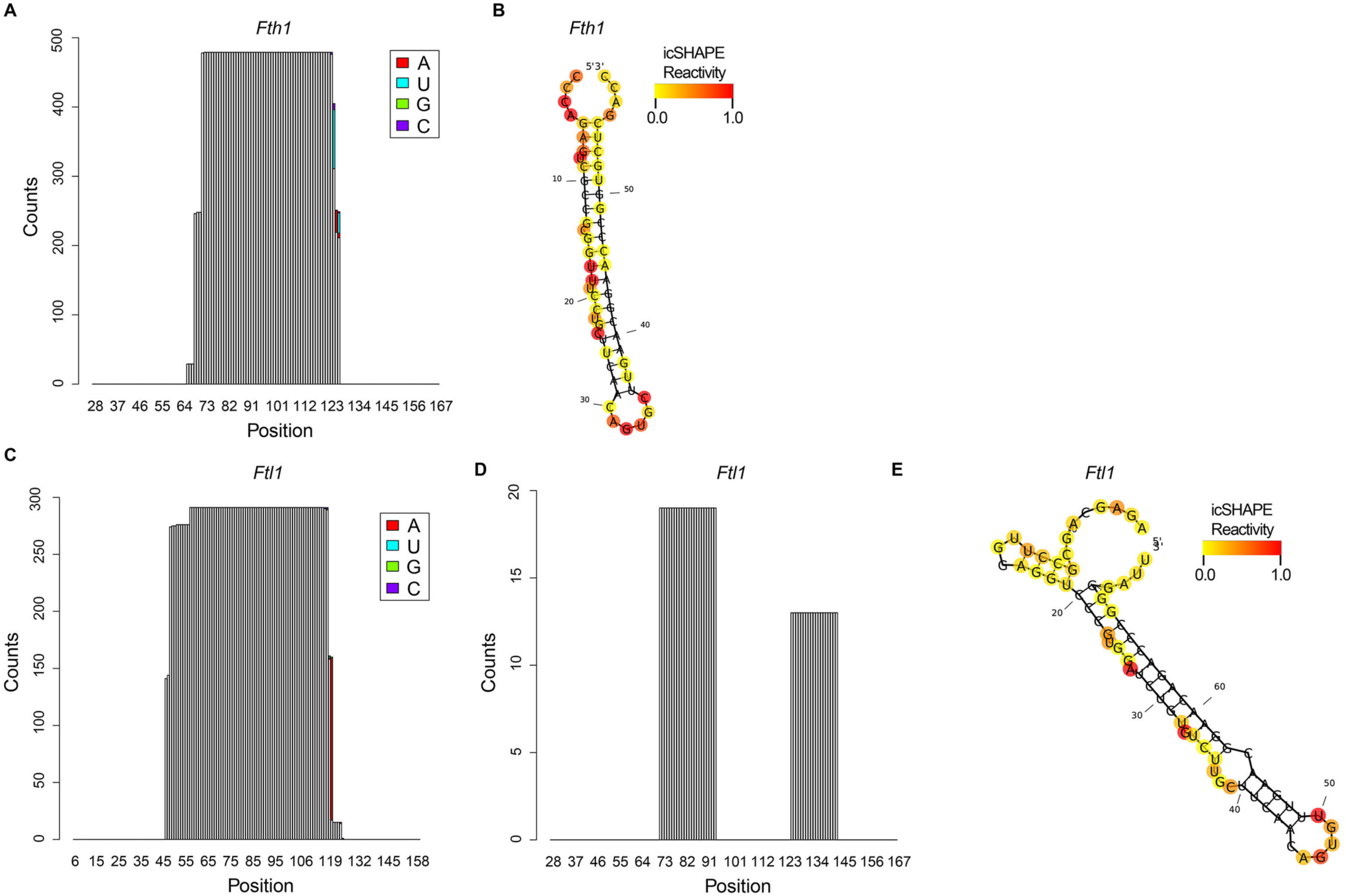

